# Root foraging response to gradients of calcium and magnesium, essential bivalent cations with low mobility in the soil

**DOI:** 10.1101/2024.03.20.584955

**Authors:** Hana Skálová, Karolína Pánková, Pavlína Stiblíková, Filip Křivohlavý, Věroslava Hadincová, Edita Tylová, Tomáš Herben

## Abstract

Plants forage for nutrients by root proliferation in nutrient-rich patches. While foraging for nitrogen and phosphorus has been repeatedly confirmed, foraging for calcium and magnesium, which are essential for plant growth and form much more stable patches in the soil, has never been examined.

We examined preferential root placement into dolomite-limestone-rich patches in a pot experiment with 17 species, and compared it with foraging for a nitrogen, phosphorus and potassium mixture (NPK). About one half of the species showed root proliferation in dolomite-rich patches. It was less pronounced than foraging for NPK and did not show any relationship to species field preferences to soil reaction, or dicots-grass difference, but it showed clear negative relationship to species-specific Ca+Mg tissue concentrations.

While foraging for NPK shows the potential of species to change their root systems by proliferation, only some species use this potential to respond to the Ca+Mg gradient. The negative correlation of this response to Ca+Mg tissue concentrations implies that nonresponding species compensate for it by physiological mechanisms. The response to Ca+Mg also implies that in contrast to nitrogen, which never shows stable patches in the soil, Ca+Mg-rich patches, which are much more stable, can be exploited by root proliferation.

## Introduction

Mineral nutrition of plants depends critically on the ability of roots to acquire nutrients from the soil. Roots are extremely plastic in response to environmental signals, by altering both their physiology and morphology. While the physiological response takes place at short temporal scales and is easily reversible (Jackson *et al*., 1990; Hutchings & De Kroon, 1994), the morphological response is different. It takes place by root proliferation (growth and branching) in soil patches rich in nutrients, thus increasing their surface and consequently uptake of nutrients in such patches (Drew, 1975; Campbell *et al*., 1991, Kembel & Cahill, 2005; Kembel *et al*., 2008). The ability to forage for nutrients by proliferation in nutrient-rich patches has been firmly established in many plant species of different habitats, growth forms and taxonomic groups (Cahill & McNickle, 2011; Chen *et al*., 2016; Wang *et al*., 2022; Stiblíková *et al*., 2023) and the physiological background recognized (Remans *et al*., 2006; Liu *et al*., 2008; Lima *et al*., 2010; Li *et al*., 2017).

Many experiments, including the original ones by Drew, 1975) were focused on individual nutrients and performed in very artificial conditions. On the other hand, the recent investigation turned to conditions closer the filed situation using solid substrates and nutrient mixtures being based on the assumption that concentrations of nutrients in the soil are correlated (Belter, 2014; Keser *et al*., 2014; Keser *et al*., 2015; Weiser *et al*., 2016). This is very different from the conditions in the field, as individual nutrients differ in their mobilities in the soil, are not consumed by plants equally and their sources are not distributed homogeneously. Thus, nutrients in the soil do not necessarily form patches where all nutrients would be present in stable ratios over time. Existing data show that temporal component of variation in nutrient concentrations at fine scales indeed strongly differs among individual nutrients and often overwhelms any spatial differences, implying that patches of individual nutrients are not or only weakly correlated (Farley & Fitter, 1999; Březina *et al*., 2019; Skálová *et al*., 2023).

The pot-based foraging experiments thus offer the plants conditions that are found nowhere in the field. The generally strong response of plant roots in the pot experiments can thus be attributed, for example, to foraging for nitrogen, which is one of the most limiting nutrients for plant growth, but almost never forms stable patches due to its fast turnover in the soil and high mobility of nitrate ions (Farley & Fitter, 1999; Březina *et al*., 2019). In contrast, less mobile elements, namely bivalent metal cations, calcium and magnesium, do show fine-scale patchy structures in the field that are more stable over time compared to patches of nitrogen (Skálová *et al*., 2023), but they have been almost completely ignored in foraging experiments in spite of their key physiological functions (Marschner, 1995) and limited availability in many soil types, namely acidic ones.

While fitness effects of root proliferation in response to nutrients are modified by a number of factors (presence of neighbours; Cahill *et al*., 2010; McNickle *et al*., 2016; Ljubotina & Cahill 2019, or limitation by other nutrients; Ward *et al*., 2008; Medici *et al*., 2015, Roger & Benfey, 2015), they critically depend on duration of nutrient patches in the soil. If nutrient-rich patches are reasonably stable, the plant will benefit from root proliferation in the patch by uptaking reasonable amounts of nutrients over the lifetime of the patch; on the other hand, if patches are short-lived, the energetical costs of building new root biomass may not be outweighed by the nutrient uptake from a fast disappearing nutrient patch (Hodge *et al*., 1999; Hodge, 2004; McNickle & Cahill, 2009; McNickle & Brown, 2014). The calcium and magnesium patches satisfy the condition of sufficient duration that may favour active proliferation in their patches, but there is no information on whether species found in soils limited by these bivalent cations possess such ability.

Compared to nitrogen and phosphorus, these cations have been paid much less attention in ecology, but they have emerged recently as an independent set of nutrients affecting a number of processes in plant life or ecosystem functions (including herbivory, palatability, decomposition rate and nutrient cycling; see e.g. Broadley *et al*., 2004; Groffman *et al*., 2011; Han *et al*., 2017, Mládková *et al*., 2018). They can be limiting in acidic soils (Marschner, 1995) and it is thus likely that ability to forage for them will provide fitness advantage. The foraging ability may boost delivery of calcium under the absence of its high-affinity influx transporters (White & Broadley, 2003). Such transporters provide the majority of uptake of other essential nutrients in roots under deficient conditions and are typically upregulated under such conditions to increase their uptake efficiency (Marschner, 1995). In contrast, we have only very scarce information about the ability of roots to modify the roots systems in response to different calcium concentrations under experimental conditions (Veatch-Blohm *et al*., 2023), and we do not know anything about roots proliferation in calcium- and magnesium-rich patches. Such ability may be ecologically relevant not only due to the low mobility of these actions, but also due to root biomass accumulation in their patches (Skálová *et al*., 2023).

We therefore examined root proliferation response to these two elements. We cultivated individuals of sixteen plant species in heterogeneous substrate with a design similar to standard foraging experiments (Campbell & Grime, 1989), but with heterogeneity only in availability of calcium and magnesium with all the remaining nutrients provided in a homogeneous fashion. We compared the magnitude of response to such heterogeneity with the response of the same species to heterogeneity in usual nutrient mixture of both forms of nitrogen, phosphate, potassium and trace elements, and compared both responses to interspecific differences in tissue concentrations of calcium and magnesium.

## Material and Methods

### The experiment

Foraging abilities were determined in sixteen species, five grasses and one graminoid (*Agrostis capillaris, Anthoxanthum odoratum, Festuca rubra, Nardus stricta, Trisetum flavescens* and *Luzula multiflora*) hereafter called graces, and 10 herbs (*Achillea millefolium, Alchemilla monticola*, *Crepis mollis, Hypericum maculatum, Lathyrus pratensis, Leontodon hispidus, Plantago lanceolata, Ranunculus acris, Trifolium repens, T. pratense*). These species are referred to by their generic names, except for the two *Trifolium* species. The species used in our study differ in their optimum soil reaction expressed by Ellenberg value for soil reaction (taken from Chytrý *et al*., 2018), which is known to correlate well with soil calcium content of their typical habitats (Schaffers & Sýkora, 2000). They range from species restricted to highly acidic and calcium-poor soils (*Nardus*) through several indifferent species to species of neutral pH and medium calcium content in the soil (*T. repens* and *Leontodon*; for Ellenberg values of all these species see Table S1 in the electronic supplementary material).

Seeds of these species were collected at a low productivity managed grassland site in the Krkonoše Mountains, North Bohemia, Czech Republic (3.75 km ESE of the centre of Pec pod Sněžkou, lat. 50°41025.165′′ N, long. 15°47041.525′′ E, 895 m above sea level). It has a cold climate (mean yearly temperature, 5.3 °C (±0.7°C); total yearly precipitation, 1275 mm; duration of snow cover, 133.6 ± 23 days (means and SDs) from the climate station at Pec pod Sněžkou from 1962 to 2020); the soil type is eutric cambisol. Soil is shallow, with very few roots extending beyond 12 cm.

The seeds were collected in July and August 2021. They were dry stored and in November placed on pure silica sand with minimum calcium and magnesium content 25.46 mg.kg^-1^ Ca and 2.48 mg.kg^-1^ Mg extracted by Mehlich III, pH 2 and measured using AAS Spectrometer ContrAA 700, Analytik Jena), slightly covered with sand and supplied by demineralized water. The germinating seeds were planted into plastic trays with individual 21 mL wells filled with sand in December and supplied with 20% Knop solution with Ca and Mg compounds replaced by ammonium and potassium ones providing the same nitrogen and sulphur supply (Table S2). In early February the plants with about six leaves, 7 cm in height and well-rooted were planted into test pots. Seeds of *Alchemilla* did not germinate until the second half of May, and thus the seedlings were transplanted into trays in early June and into the test pots only in late June; attention was paid to used seedlings of comparable size to those used in February.

The test pots were round 3-L pots (inner top diameter 19 cm, bottom diameter 16 cm, TEKU Pöppelman, MCI 19). The test pots were filled with silica sand (using a non-woven fabric at the bottom of the pot to prevent the sand from falling out and limit the root growth outside the pot) with a cylindrical patch with 2 cm in diameter at a pot side filled with a 12 g of fine-grained dolomitic limestone, containing 190 g.kg^-1^ Ca and 11 g.kg^-1^ Mg, mixed with the sand to get the volume of 40 ml. Even if using dolomitic limestone disable disentangling foraging for individual elements, we are using it as soil concentrations of these two elements are highly correlated and therefore it constitutes a meaningful ecological signal as such (Skálová *et al*., 2023). The patch was created by a specially designed instrument that ensured vertical position of the patch and the distance of 5 cm from the pot centre and 1 cm from the pot side at the bottom to enable the harvest of the patch and its close surrounding (see Fig. 1). The amount of dolomitic limestone needed to create a sufficiently strong gradient across the pot was determined in a previous pilot experiment.

**Fig. 1:**
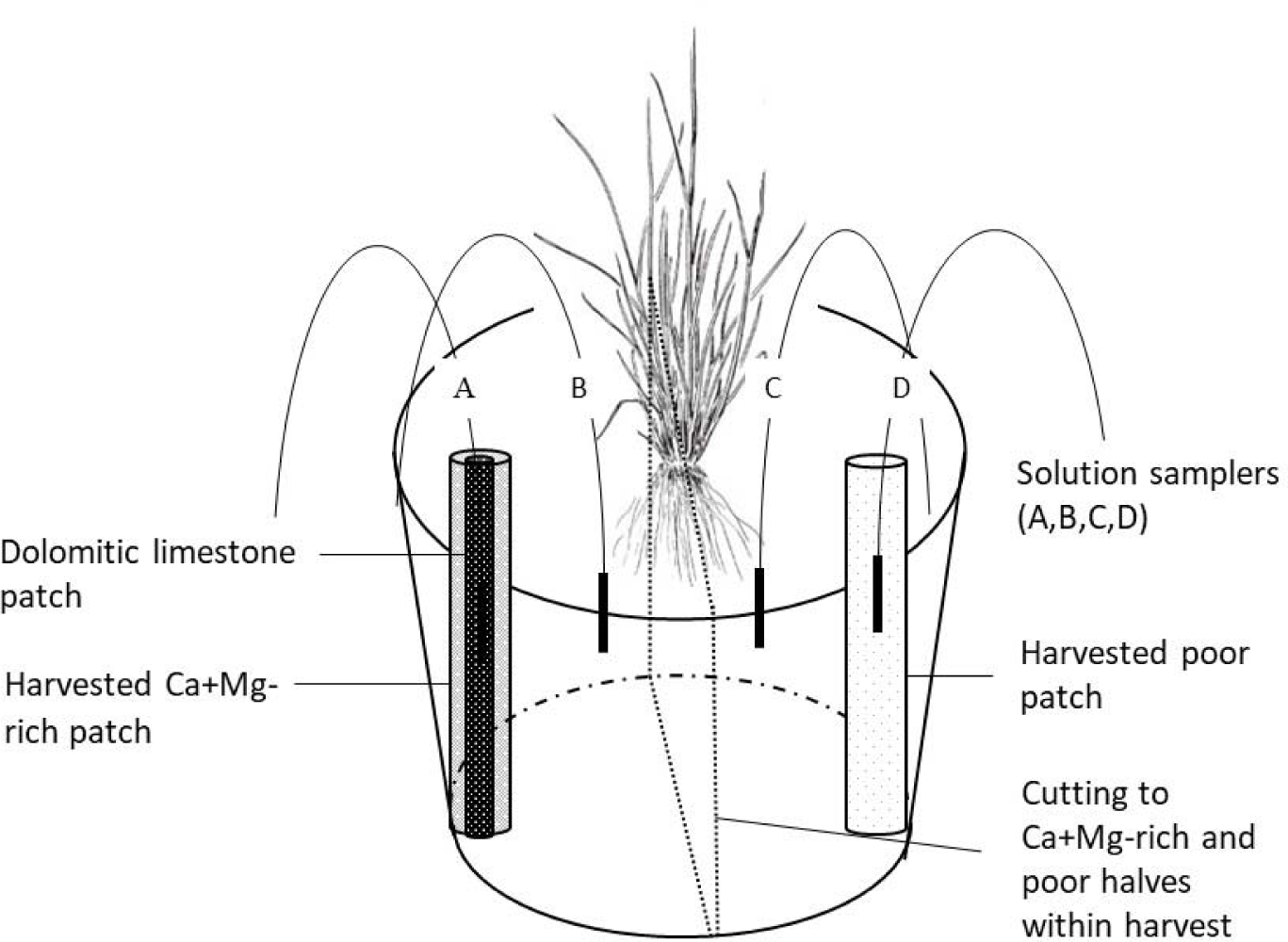
Spatial arrangement of plant, dolomitic limestone patch and sampler positions and individual harvested parts within the pot.

To check whether Ca and Mg concentrations indeed form gradients as expected, we sampled the solution present in the sand three times during the experiment (in early March, mid-April and before the harvest in May). The sampling was done using Rhizon SMS samplers with a diameter of 2.5 mm, length of 10 mm and mean pore size of 0.15 µm (Rhizosphere Research Products B.V., The Netherlands, https://www.rhizosphere.com) with 60 ml syringes with 5 cm wooden retainers attached to the samplers to create sufficient vacuum 1068 Pa). The samplers were installed in three pots per species. In each pot there were four samplers placed at 6 cm distances along a line across the pot in the depth of 5 cm. One sampler was at the place of the dolomite patch, one at the opposite site of the pot and two samplers equidistantly between them, i.e. closer to the pot centre (Fig. 1). Ca and Mg contents in the solutions were measured using Agilent 5900 SVDV ICP-OES, wavelengths (Ca: 393,366 + 396,847 + 422,637 nm; Mg: 279,553 + 280,270 nm). Samples were diluted due to different volume in range of 1:0 to 1:100 and measured in polypropylene vials. These measurements showed that Ca and Mg concentration in the soil solution within the patch were about 10-times larger than the average in the soil solution collected from the depth of 5 cm in the community where the seeds came from (Skálová et al. 2023; compare Table S3 and Fig. S1). Values at the opposite side of the pot (with no dolomitic limestone) were close to the field values for Ca, and lower than field values for Mg. The average concentration of the elements in the soil solution decreased during the experiment while the contrast between the sampler placed in the patch and the remaining samplers increased, most likely due gradual release of the elements from the patch and also due to absorption of both elements by plant roots (Fig. S1). The decline over time was highly statistically significant for both elements, but it did not differ among the species (Table S1).

Individual plants were planted in centres of these test pots in 13 replicates per species. Once or twice a week, depending on the temperature and plant size, the pots were supplied from the bottom with the same 20% Knop solution as the seedlings while the pots were supplied till full soaking capacity. In mid-April when temperature and evapotranspiration considerably increased, we switched to 10% Knop solution provided twice a week. Similar pattern was used in *Alchemilla* planted later than the other species*. Alchemilla* planted in June was supplied 10% Knop solution till mid-September and then by 20% Knop solution to compensate for the different course of evaporation during the growth period of this later set of pots.

The seeds were germinated and plants cultivated in a greenhouse with temperature regulation; thus, the temperature was about 2-7°C higher than outside, but 10°C and 15°C at minimum at night and during the day, respectively (for monthly outside temperatures during the cultivation time see Table S4). In March, additional light was provided between 7 AM and 7 PM by fluorescent lamps Osram Fluora L36/77 placed about 1 m above the pots.

The plants were harvested 14-15 weeks after planting to the cultivation pots (see Fig. S2). In *Alchemilla*, attention was paid to harvest plant comparable with the other species harvested in the first run. The aboveground biomass was cut at the substrate level, dried for ten hours at 60°C and weighed. The patch was excavated using a tube of inner diameter of 3 cm attached to the modified instrument used for its creation. It enabled us to excavate the patch 2 cm in diameter) plus additional layer of soil around it of 0.5 cm where high concentration of roots can be expected in case of foraging species; hereafter called Ca+Mg-rich patch, see Fig. 1. An identical 3 cm diameter sample was taken from the opposite side of the pot at the same distance from the rooting plant as a control; hereafter called poor patch. After that each pot was divided into two halves in the middle of the plant’s rooting point by a sharpened iron sheet with the Ca+Mg-rich patch situated perpendicular to the cut in the maximum distance from the cut. Each half was weighed for the later possible correction of root density. The half of the pot containing the Ca+Mg-rich patch is hereafter called Ca+Mg-rich half and that with poor patch is called poor half. The roots that were not possible to assign to any of the four above samples (roots attached and overgrown in the non-woven fabric at the bottom of the pot) were harvested separately and included in the total root biomass only. For roots, all five samples (rich and poor patches, rich and poor halves of the pot and remaining unassigned roots) were washed in water on a fine sieve (0.8 mm). The plant bases and rhizomes were separated, dried and weighed and included into the aboveground biomass.

### Species-specific tissue concentrations of calcium and magnesium

Calcium and magnesium content was measured in three separate tissues: (i) the aboveground parts of the plants harvested in the experiment (excluding bases and rhizomes), and (ii) roots from the pot poor halves. We used material from three randomly chosen individuals per species. Nonflowering shoots were taken for grasses and graminoids and leaves for dicotyledons except *Crepis* where fertile shoots were used to get enough material for analyses. Further, (iii) we sampled aboveground tissues of all species in the second half of June 2022 (i.e., at the top of the vegetation season) at the same locality (within an area of approximately 50 x 50 m) where the seeds had been collected. Plant parts comparable with that from the experiment were taken. We refer to these three sources of tissues for the analysis (aboveground biomass from the experiment, roots from the control part of the pot in the experiment, and aboveground biomass from the field) as sample types. The samples were ground (to a particle diameter < 0.1 mm), dried at 60 °C and mineralized (0.5 g plant biomass + 4 ml HNO_3_ +2 ml 30% H_2_O_2_) in the Milestone Standard 1200 Mega mineralization unit. Ca and Mg concentrations were measured by absorption atomic spectrophotometry in H_2_SO_4_ and LaCl_3_ to eliminate the effects of phosphates and sulphates (Moore and Chaoman 1986) using AAS Spectrometer ContrAA 700, Analytik Jena, with C_2_H_2_-air flame for Mg and C_2_H_2_-N_2_O flame for Ca.

As there were strong correlations of Ca and Mg content in tissues of individual species across individual sample types and elements (Table S3 and Fig. S3), we summarized overall concentrations of both elements in tissues of individual species by taking scores of these species on the first axis of principal components analysis of standardized values of aboveground Ca and Mg concentrations (Fig. S3). This axis accounted for 71% of total interspecific variation in Ca and Mg. Principal components analysis was calculated using the function rda from the package vegan (Oksanen *et al*., 2020, version 2.5-7). Tests of difference between grasses and forbs in their Ca and Mg content were done using mixed model analysis of variance with species as a random factor, fitted using the lmer function from the lme4 package and tested by Type III sum of squares analysis of variance with Satterthwaite’s approximation of denominator degrees of freedom (function anova.lmer from the lmerTest package, version 3.1-3, Kuznetsova *et al*., 2017).

### Data analysis

We used two measures of root foraging: (i) differences in root density between Ca+Mg-rich patch and the identically sized poor sample taken at the opposite side of the pot ("patch-level foraging"), and (ii) between the Ca+Mg-rich pot half and the poor half ("pot-level foraging"). Both these differences were examined by standard major axis regressions of the root biomass in the Ca+Mg-rich part against root biomass in the poor part, forced through origin. We tested whether the slope of such regression line significantly differs from one (a value expected under the null hypothesis of equal root placement into the Ca+Mg-rich and poor patches). Root biomasses were square-root transformed to improve normality of errors and the slope estimates were back-transformed to linear scale. Tests were run for each species separately. In addition, we performed a global test to determine whether slopes are significantly different among the species by fitting a model of the root biomass in the Ca+Mg-rich part against species and root biomass in the poor part at the opposite side of the pot, and testing interaction between the two predictors by log-likelihood ratio test. The standard major axis regressions were fitted using the R package smatr (Warton *et al*., 2012). All statistical calculations were done in the R environment (version 4.1.3; R Core Team 2022).

Differences in foraging between grasses and dicots were tested using mixed model analysis of variance with species as a random factor. Tests were done with Type III sum of squares and Satterthwaite’s approximation of denominator degrees of freedom. Models were fitted using lmer function from the lme4 package (version 1.1-27.1, Bates *et al*., 2015) and tested by lmerTest package (version 3.1-3, Kuznetsova *et al*., 2017).

Data on root proliferation for in pots with gradients of nutrient mixture (NPK) were taken from Herben *et al*. (2022) and Stiblíková *et al*. (2023) where 14 species tested for Ca+Mg foraging were included (i.e. *Alchemilla*, *Hypericum* and *Trisetum* were missing in these studies). In these studies, plant individuals were exposed to heterogeneous pot treatment generated by two drippers positioned at opposite ends of the pots, one with water and the other with commercial fertilizer (0.2% dilution of Wuxal Super NPK 8 : 8 : 6 + micronutrients, but no Ca and Mg; Aglukon). For consistency with the current foraging calculation, foraging was expressed in the same way as in the current experiment (i.e. slope of the standard major axis regression forced through origin after square-root transformation of root biomasses). For detailed description of the design, see Herben *et al*. (2022) and Stiblíková *et al*. (2023). We further expressed specific Ca+Mg foraging as the difference between log-transformed foraging for NPK and log-transformed foraging for Ca+Mg; negative values thus mean that Ca+Mg foraging in the given species is weaker relative to the NPK foraging.

## Results

Species significantly differed in their ability to place roots to the calcium and magnesium rich patches (likelihood ratio test of the interaction between species and pot part: 53.26, d.f = 15, P<0.001) as well as in the ability to place roots in the Ca+Mg-rich halves of the pot (likelihood ratio test of the interaction 93.19, d.f = 15, P<0.001). The species with the most consistent ability to forage for Ca+Mg were *Ranunculus, Lathyrus, Agrostis* and *Anthoxanthum* (Table 1, Fig. 2, 3, and Fig. S3), although the numerical values of preferential root placement strongly differed among species. *Ranunculus* placed into the Ca+Mg-rich patches more than four times as many roots than into the corresponding poor patches; this ratio was 2.52, 1.93 and 1.58 in *Lathyrus, Agrostis* and *Anthoxanthum*, respectively. The ratios at the pot-level were always much smaller, with *Ranunculus* having the highest ratio of 1.63, and the other three species around 1.3 (Fig. 3). Three species avoided Ca+Mg-rich halves of the pot (*Luzula*, *Hypericum*, and *Alchemilla*); but no species avoided the Ca+Mg-rich patch. There was no significant difference between grasses and forbs in Ca+Mg foraging either at the patch or pot level (patch level: F = 0.045, d.f. = 1 and 13.7, P = 0.835; pot level: F = 0.821, d.f. = 1 and 15.1, P = 0.379).

**Fig. 2:**
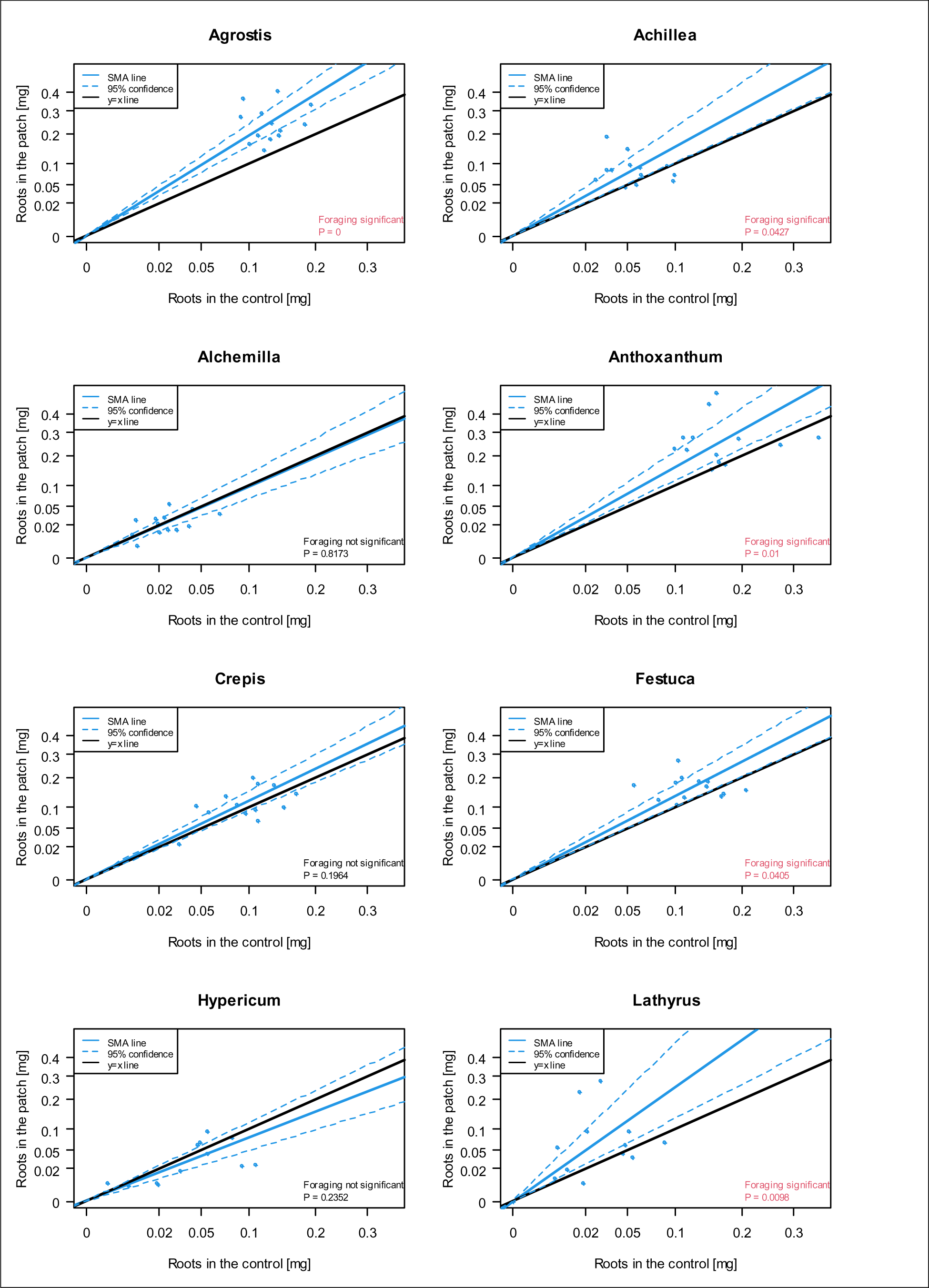
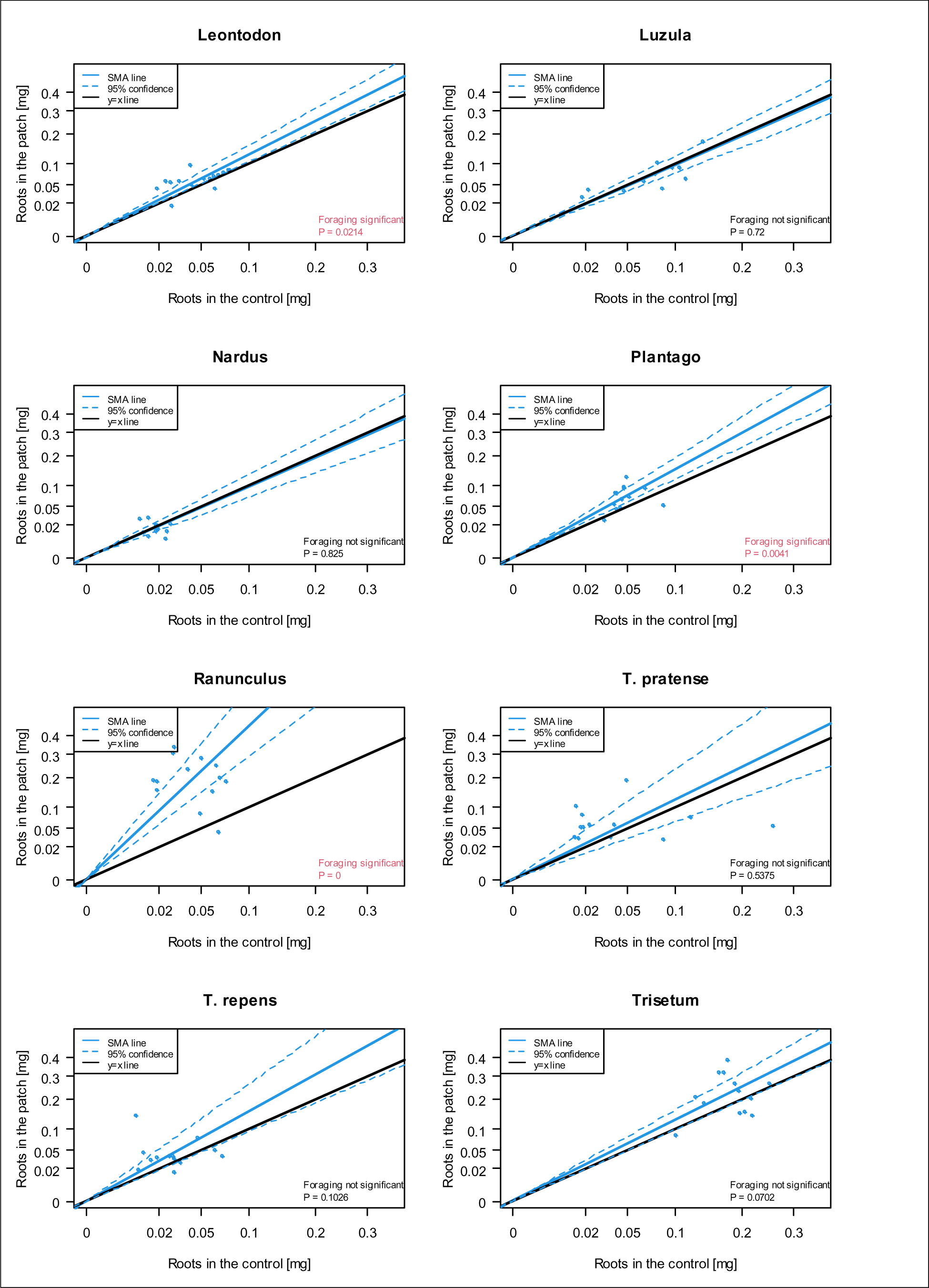
Root placement into rich and poor patches in individual species. Each point indicates one pot. The lines were fitted using standard major axis regression forced through origin. The blue line is the standard major axis regression line, the dashed lines show 95% confidence intervals for the slope parameter; the black line is the y=x line (expectation under the assumption of no preferential placement). Foraging is considered significant if the slope is greater than one and the 95% confidence interval of it does not cover one. Note the square root scaling of the axes. For a similar figure showing differences between pot halves see Fig. S3.

**Fig. 3.**
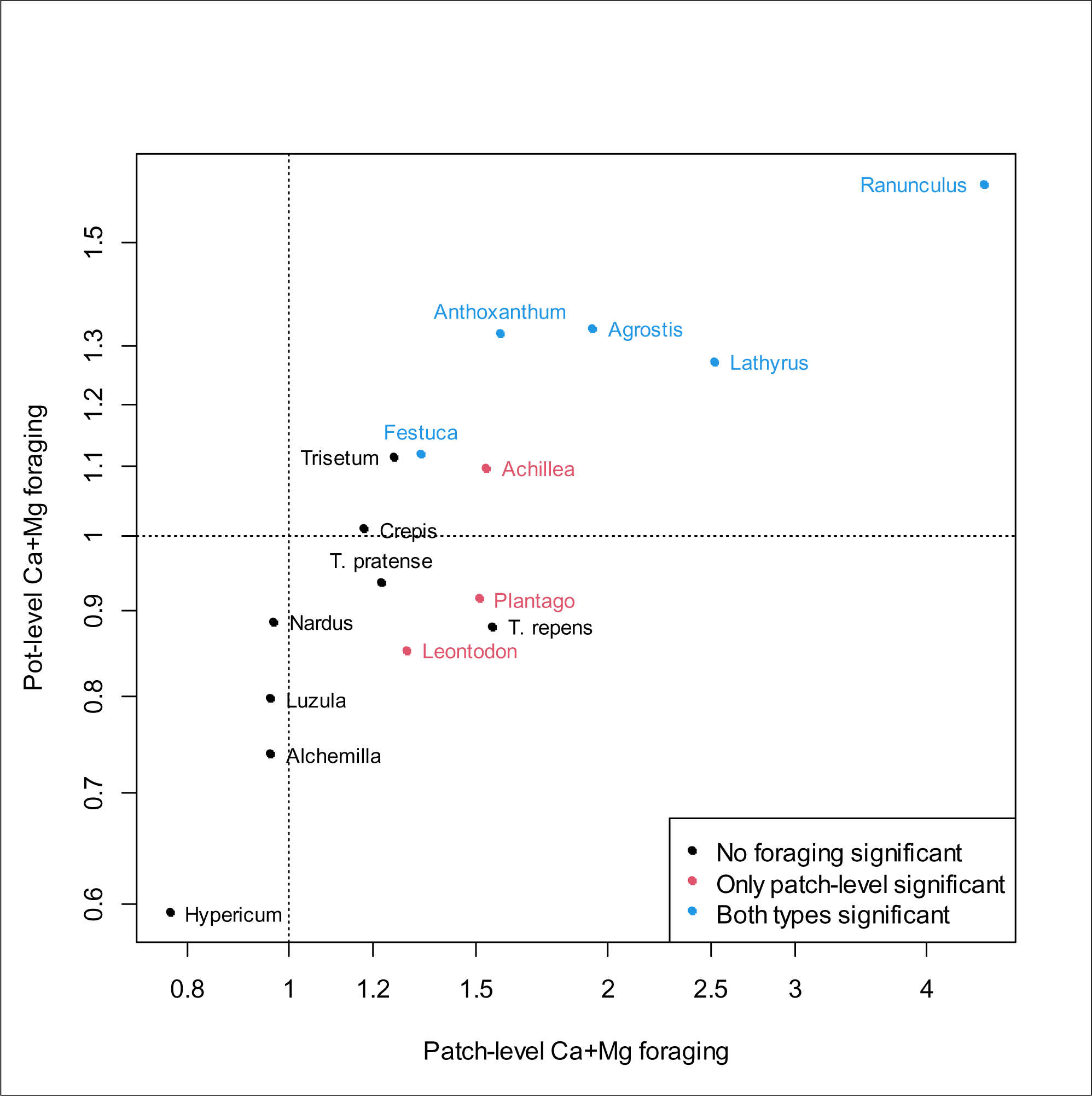
Relationship between patch-level and plot-level foraging. Foraging is expressed as a slope of the relationship between root biomass in the rich and poor parts (see the Methods for details). Note the log-scaling of the axes; value of one thus means no foraging. For data on individual pots see Fig. 2, and the Fig. S3.

**Table 1.**
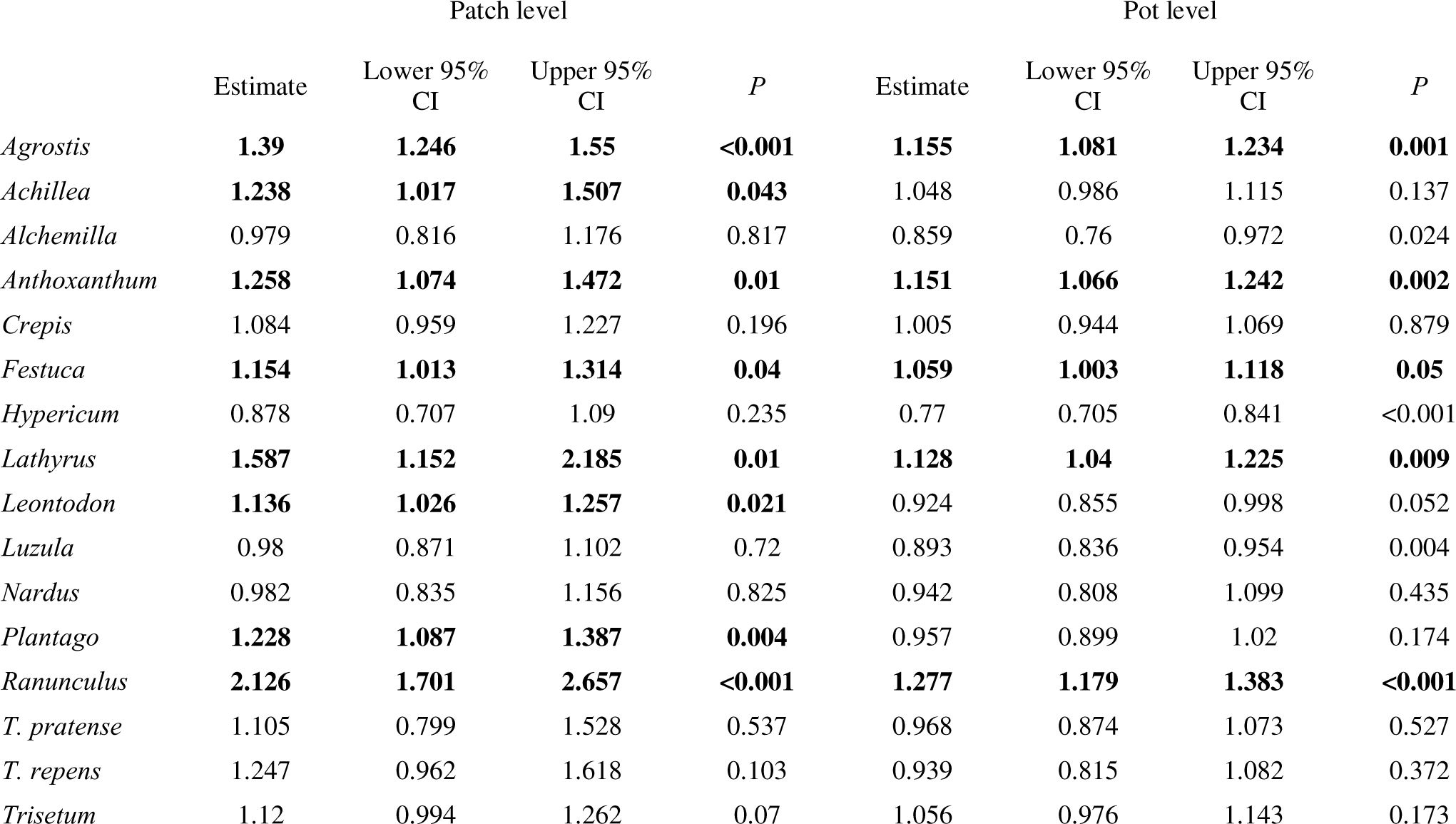
Best estimates of slopes of relationship between rich and poor parts in the pot. Slopes larger than one are indicative of foraging, i.e. preferential placement of roots into the calcium and magnesium rich part. Species with significant values larger than one are shown in bold. *P* is significance of the two-sided test of difference from unity (the expectation under no foraging). For plots of individual species, see Figure 2 and Fig. PLOTS-SPECIES in the electronic appendix.

|  | Patch level |  |  |  | Pot level |  |  |  |
| --- | --- | --- | --- | --- | --- | --- | --- | --- |
|  | Estimate | Lower 95% CI | Upper 95% CI | *P* | Estimate | Lower 95% CI | Upper 95% CI | *P* |
| *Agrostis* | **1.39** | **1.246** | **1.55** | **<0.001** | **1.155** | **1.081** | **1.234** | **0.001** |
| *Achillea* | **1.238** | **1.017** | **1.507** | **0.043** | 1.048 | 0.986 | 1.115 | 0.137 |
| *Alchemilla* | 0.979 | 0.816 | 1.176 | 0.817 | 0.859 | 0.76 | 0.972 | 0.024 |
| *Anthoxanthum* | **1.258** | **1.074** | **1.472** | **0.01** | **1.151** | **1.066** | **1.242** | **0.002** |
| *Crepis* | 1.084 | 0.959 | 1.227 | 0.196 | 1.005 | 0.944 | 1.069 | 0.879 |
| *Festuca* | **1.154** | **1.013** | **1.314** | **0.04** | **1.059** | **1.003** | **1.118** | **0.05** |
| *Hypericum* | 0.878 | 0.707 | 1.09 | 0.235 | 0.77 | 0.705 | 0.841 | <0.001 |
| *Lathyrus* | **1.587** | **1.152** | **2.185** | **0.01** | **1.128** | **1.04** | **1.225** | **0.009** |
| *Leontodon* | **1.136** | **1.026** | **1.257** | **0.021** | 0.924 | 0.855 | 0.998 | 0.052 |
| *Luzula* | 0.98 | 0.871 | 1.102 | 0.72 | 0.893 | 0.836 | 0.954 | 0.004 |
| *Nardus* | 0.982 | 0.835 | 1.156 | 0.825 | 0.942 | 0.808 | 1.099 | 0.435 |
| *Plantago* | **1.228** | **1.087** | **1.387** | **0.004** | 0.957 | 0.899 | 1.02 | 0.174 |
| *Ranunculus* | **2.126** | **1.701** | **2.657** | **<0.001** | **1.277** | **1.179** | **1.383** | **<0.001** |
| *T. pratense* | 1.105 | 0.799 | 1.528 | 0.537 | 0.968 | 0.874 | 1.073 | 0.527 |
| *T. repens* | 1.247 | 0.962 | 1.618 | 0.103 | 0.939 | 0.815 | 1.082 | 0.372 |
| *Trisetum* | 1.12 | 0.994 | 1.262 | 0.07 | 1.056 | 0.976 | 1.143 | 0.173 |

At the level of individual species, there was a fairly strong relationship between patch-level and pot-level foraging (Fig. 3). In most species, the patch-level foraging was stronger compared to the pot-level foraging and was more often significant; *Achillea, Plantago* and *Leontodon* showed significant patch-level foraging, but not pot-level foraging (Table 1, Fig. 3). There was no significant difference between grass and forb species in either type of foraging (analysis of variance, d.f. = 1, 15) nor was there any significant relationship to the species optimum of soil reaction expressed by the Ellenberg indicator value of soil reaction, which is known to capture largely Ca soil content in their optimum habitats (linear regression, d.f. = 1, 15).

Relationships between Ca+Mg foraging and foraging for NPK were not significant (linear regression, F = 0.3911 on 1 and 12 d.f., P = 0.5434, and F = 1.936 on 1 and 12 d.f., P = 0.1893, for patch- and pot-level foraging respectively, see Fig. 4), but the relationship was primarily triangular. While some species had similar values of foraging for Ca+Mg-rich and NPK (*Anthoxanthum, Lathyrus, Agrostis* among strong foragers, and *Crepis, Nardus* and *Festuca* among weak foragers), there was a group of species which were able to forage for NPK, but did not respond to Ca+Mg (namely *Leontodon,* both *Trifolium* species, *Plantago*). Only *Ranunculus* was a better forager for Ca+Mg compared to NPK (Fig. 4).

**Fig. 4:**
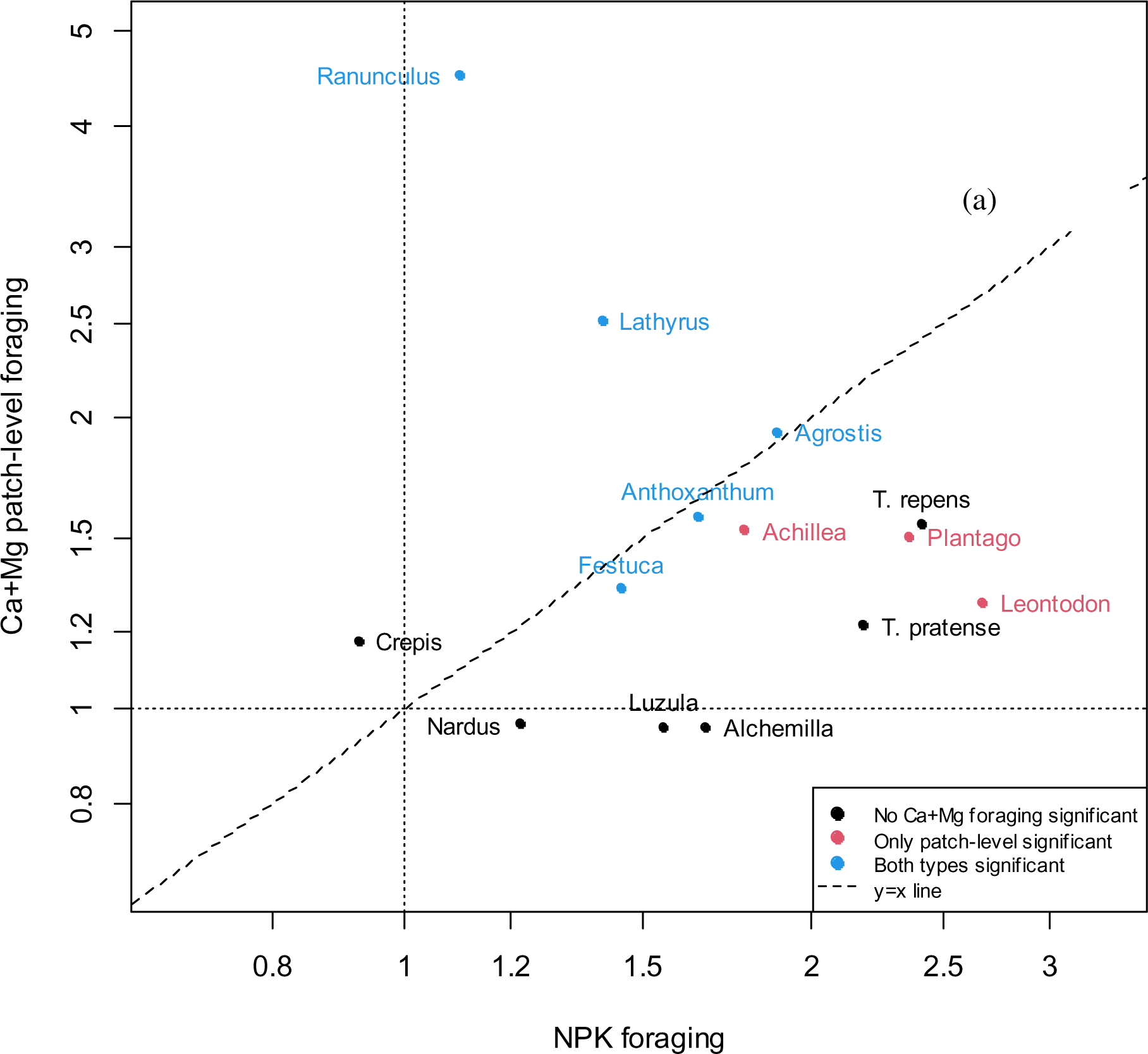
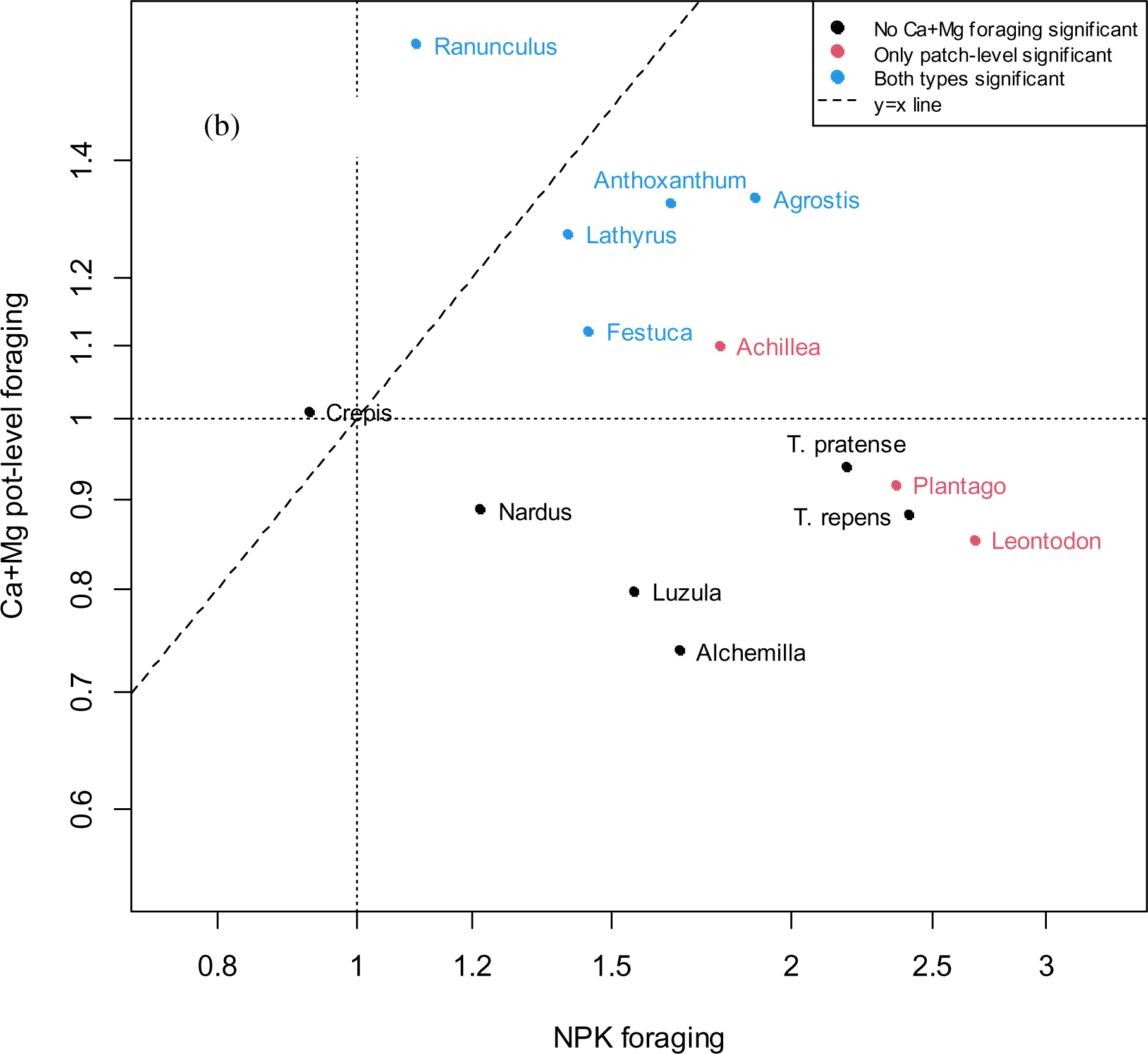
Relationship between calcium and magnesium foraging and NPK foraging (foraging for NPK + essential nutrients). Foraging is expressed as slope of the relationship between root biomass in the rich and poor parts. Note the log-scaling of the axes. Patch-level foraging (a) is determined at the level of Ca+Mg-rich cylindrical patches of 2 cm in diameter; pot-level foraging is determined at the level of whole calcium and magnesium rich and poor halves of the pot.

There were strong and significant differences in Ca+Mg tissue concentrations among species and among sample types, with concentrations in aboveground tissues being markedly larger than those in roots and concentrations of Mg (but not of Ca) being higher in the field compared to the experiment. In both elements, there was strong interaction between species and sample type, indicating that individual species differed both in root-shoot and experiment-field comparisons (Fig. 5, Table S6 in the electronic supplementary material). Forb tissues had significantly higher concentration for both Ca and Mg than grasses (using mixed model anova with species as a random factor, Table S6, Fig. S5). There were strong correlations of Ca and Mg content in tissues of individual species across individual sample types and elements (Table S3). Consequently, in principal components analysis of standardized values of six variables (Ca and Mg concentrations in three sample types), the first axis accounted for 71% of the total variation, indicating high degree of intercorrelation of both elements in plant tissues irrespective of the sample type (Fig. S4).

**Fig. 5:**
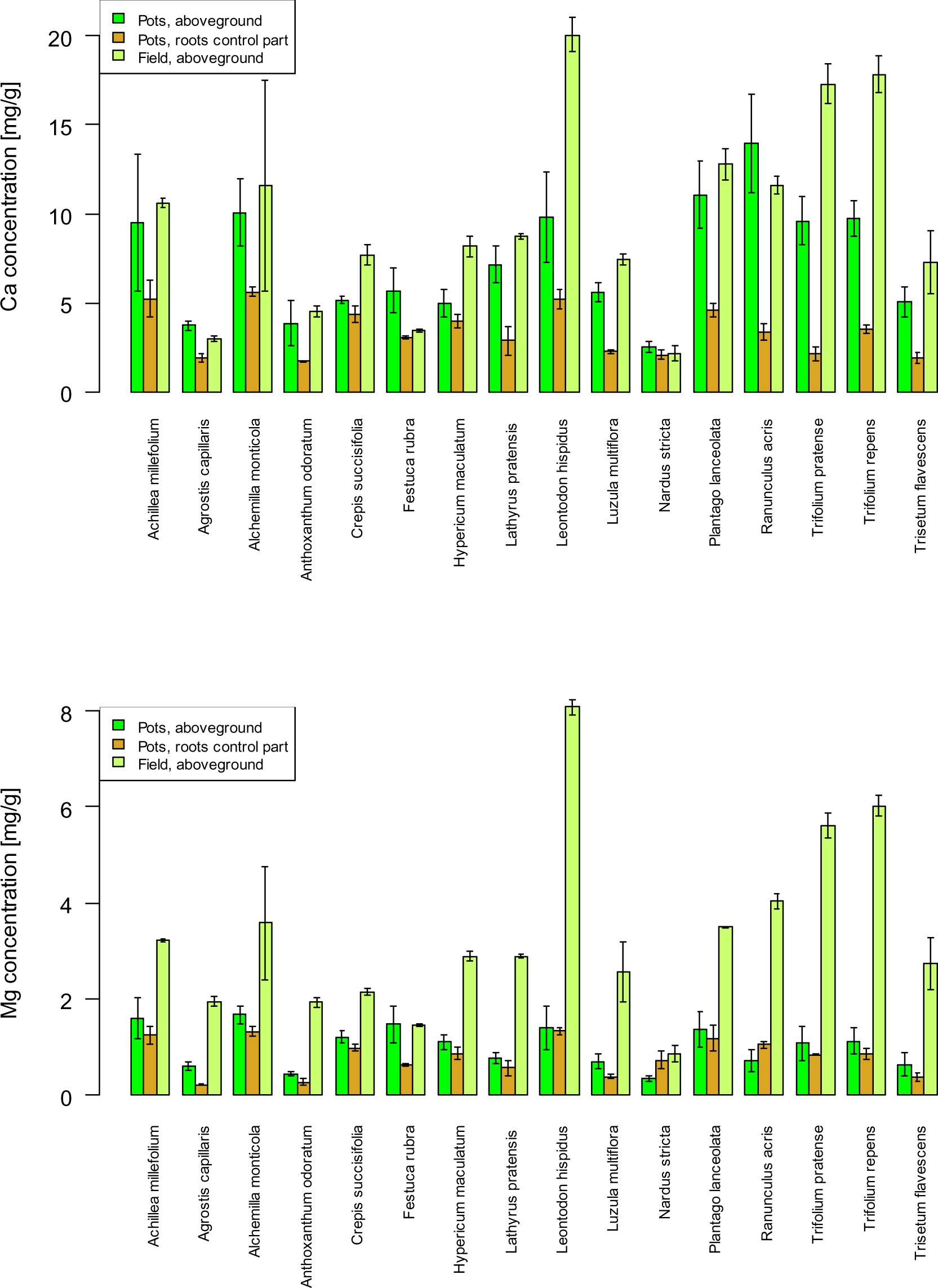
Concentration of calcium and magnesium in plant tissues of individual species. Error bars are 95% confidence intervals.

There was a weak negative relationship between pot-level foraging for Ca+Mg and concentration of these elements in plant tissues (Fig. S5). When foraging for Ca+Mg was expressed as a difference from foraging for NPK, it showed strong and significant negative relationship to shoot concentrations of Ca and Mg, which was even stronger for dicot species only (Table 2, Fig. 6, Fig. S6 in the electronic supplementary material).

**Table 2.** Tests of the effect of calcium and magnesium shoot concentrations and grass/nongrass on the difference between calcium and magnesium foraging and NPK foraging.

|  | d.f. | *F* | *P* |
| --- | --- | --- | --- |
| calcium and magnesium shoot concentration | 1 | 8.238 | 0.015 |
| Grass | 1 | 5.397 | 0.040 |
| Residuals | 11 |  |  |

Patch-level
|  | d.f. | *F* | *P* |
| --- | --- | --- | --- |
| calcium and magnesium shoot concentration | 1 | 2.766 | 0.125 |
| Grass | 1 | 4.880 | 0.049 |
| Residuals | 11 |  |  |

**Fig. 6.**
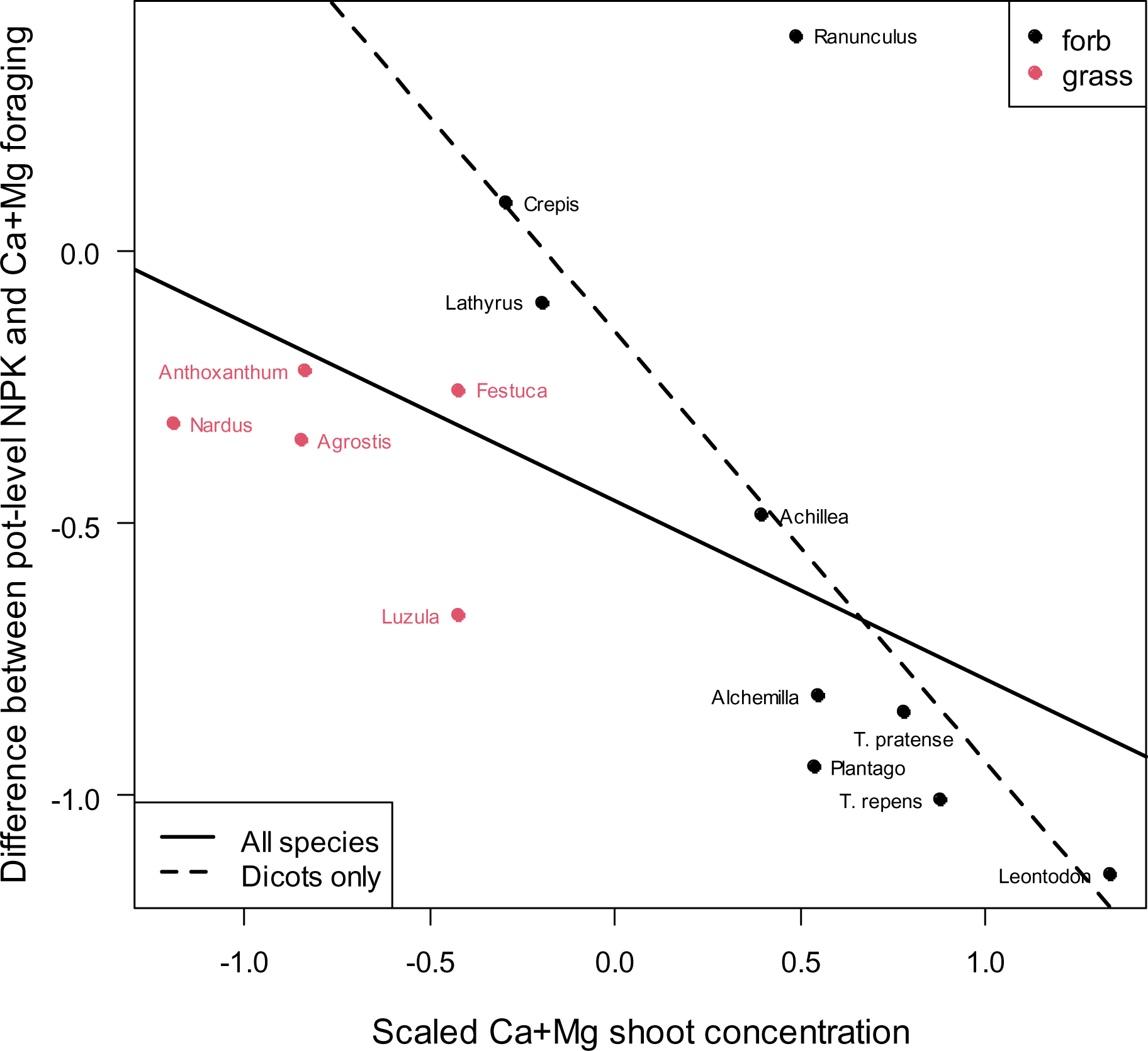
Relationship between calcium and magnesium shoot tissue concentration and foraging for calcium and magnesium. Foraging is expressed as difference between NPK foraging (pot-level) and calcium and magnesium patch-level foraging (see Fig. GENERAL); negative values thus mean that calcium and magnesium foraging in the given species is weaker relative to the NPK foraging. Calcium and magnesium shoot concentration is the concentration of both elements in aboveground shoots (both in experiment and in the field). The full line is the least-squares relationship for all species (*F* = 6.0288, d.f. = 1,11, *P* = 0.03029), the dashed line the least-squares relationship for dicots only ((*F* = 10.532, d.f. = 1,7, *P* = 0.01415). grasses – grasses and graminoids (*Luzula*).

## Discussion

The experiment clearly showed that calcium and/or magnesium gradients can elicit plastic response by the root system in a number of plant species. This foraging ability was fairly fine-scale, taking primarily place at the scale of centimetres, which is comparable with root and cation heterogeneity in the field of seed origin (Skálová *et al*., 2023) and possibly more generally (Farley & Fitter 1999). Due to their chemical properties, calcium and magnesium tend to form much more stable patches in the soil, making foraging for them a reasonable ecological strategy. Our finding also indicates that the root aggregation in Ca+Mg rich patches in the field is likely to be generated by heterogeneity in cation distribution and not vice versa, which was not clear from the field study (Skálová *et al*., 2023). However, it is important to keep in mind that root proliferation is only one component of the processes responsible for nutrient acquisition, and the other components may compensate for low root proliferation.

However, root proliferation in Ca+Mg rich patches was by no means universal across species. In a majority of the species involved in the experiment, the ability to forage for Ca+Mg was linked with their ability to forage for NPK, but there was a remarkable group of species that were able to forage strongly for NPK, but did not respond to the Ca+Mg concentration gradient. (In contrast, there was only one species, *Ranunculus*, that responded to the Ca+Mg signal more strongly compared to NPK.) This is likely to indicate that the ability to change root system shape, which is perhaps best shown by their response to often-limiting nutrients such nitrogen and phosphorus, is a necessary condition for foraging for cations. However, it may be noted here that foraging for nitrogen, despite being often studied in foraging experiments, seems to be of limited ecological relevance due to very short timespan of its patches in the field, while foraging for the elements that occur in more stable patches under field conditions like Ca and Mg is much more likely to be ecologically relevant.

Further, the ability to forage for calcium and magnesium was significantly lower in species with high tissue concentrations of these elements. High Ca tissue concentration thus does not need to reflect high demand for this element, but it may reflect high occurrence of calcium oxalate crystals in vacuoles (Paiva, 2019). Due to its function as secondary messenger, plants precisely control Ca^2+^ concentration in cytoplasm and the superfluous Ca^2+^ ions actively exported from the cytoplasm and immobilized as calcium oxalate. As Ca^2+^ also plays a structural role by crosslinking pectin molecules (Thor, 2019) lower abundance of pectin in cell walls may explain low concentrations of Ca in such tissues (Broady *et al*., 2003). Unlike other essential nutrients, Ca does not have specific high-affinity transporters and Ca is taken up mostly by root apoplast along the concentration gradient, and only temporarily enters symplast when passing endodermal layer with Casparian bands of young root (White & Broadley, 2003; Cholewa & Peterson, 2004; Thor, 2019). In addition, retranslocation within plant bodies common in other nutrients is generally limited in Ca (Paiva, 2019). These two factors may make foraging for Ca more important.

Foraging for magnesium by root proliferation is likely to be modified due to presence of specific transporters of the MGT family (Mao *et al*., 2014) and its higher phloem mobility than Ca (Broadley *et al*., 2008). However, the uptake of Mg may be strongly limited by other cations including H^+^ that can result in a fairly widespread magnesium deficiency (Marschner, 1995) and may thus make foraging reasonable. The negative relationship between pot-level foraging for Ca+Mg and concentration of these elements in plant tissues, especially in dicots, supports the idea about the compensatory effect of foraging for these elements.

Interestingly, three species (*Luzula, Hypericum, Alchemilla*) had significantly more roots in pot halves without the Ca+Mg patch, although they did not show any difference in root density at the fine scale of patches. At least two of these species come from fairly cation-poor habitats, but otherwise do not have much in common (they differ in Ca and Mg tissue concentrations, and in the ability to forage for NPK). As Ca limits acquisition of other cations including NH_4_^+^ that is important source of nitrogen in plants under acidic soil conditions, roots of these species might be presently placed into the halves without the Ca+Mg patch to ensure sufficient nitrogen supply.

When comparing the reported experiment with data on foraging NPK, it must be kept in mind that both foraging types were assessed by slightly different designs. As nitrate ions and soluble phosphorus are more mobile in soil compared to Ca and Mg, the nutrient heterogeneity must be maintained by dripping nutrient solution to yield a fairly sharp contrast between the pot halves. On the other hand, such system could not be used for creating Ca+Mg heterogeneity, as the dripping system used tap water under pressure with Ca and Mg concentration higher than the soil water from the mountain meadow where the seeds originated. This necessitated using deionized water for Ca and Mg. Gradients of less mobile Ca and magnesium Mg can then be easily maintained by adding finely ground dolomitic limestone to small patches in the pots.

Consequently, comparison of the whole pot halves only does not need to be as informative as they are in NPK foraging experiment; indeed, our results clearly show that root abundance in the close surrounding of the patch and in the corresponding sample from the opposite site of the pot provide much clearer image of species response to gradient in calcium and magnesium. It should also be noted that this approach does not permit separation of foraging response to Ca from the response to Mg. However, their concentrations are generally highly correlated in the field (see e.g., Skálová *et al*., 2023; Ritz *et al*., 2004). Therefore, joint concentration gradient of both Ca and Mg constitutes a meaningful ecological signal.

### Conclusions: ecological relevance of the findings

Our experiment demonstrated, for the first time, active root proliferation in response to concentration of bivalent cations in heterogeneous substrate. In contrast to highly mobile nitrogen ions which show high turnover in the soil, fine-scale calcium and magnesium concentration patches tend to be much more stable, making investment into root biomass associated with foraging for them rewarding for the plant. This implies that gradients of these elements may contribute to heterogeneity of root systems in the field, primarily in habitats where these elements are in short supply.

## Acknowledgements

The Krkonoše National Park Administration kindly permitted the experiments to run on their meadows (contract number OSML 38-4/2018). Our thanks are also due to a number of people who helped us with the plant harvest and washing roots. The research was funded by the GAČR grants 20-02901S and 23-05654S.

## Author contributions

TH, HS and PS designed the research. HS led the experimental part of the study, HS, KP and VH performed the experiment and FK did the chemical analysis. TH and HS analysed the data, and HS, ET and TH interpreted the analyses and wrote the text with contributions of all other authors. All authors approved the final version of the manuscript.

## Electronic supplementary material

Table S1. Ordinal values (Ellenberg-type values; Chytrý et al. 2020) expressing species preference to calcium content in the soil.

Tab S2. Composition of classical and modified 20% Knop solution (mg/L).

Table S3. Tests of differences in calcium and magnesium concentrations in experimental pots.

Table S4. Average daily temperatures during the experiment

Table S5. Correlation coefficients of calcium and magnesium content in plant biomass across individual sample types.

Table S6. Analysis of variance of calcium and magnesium content in biomass across species and across sample types

Table S7. Tests of differences in calcium and magnesium content in biomass between grasses-forbs and among sample types

Fig. S1: Calcium and magnesium concentrations in the pots during the experiment.

Fig. S2: Experimental plants in the greenhouse at the harvest time

Fig. S3 Root placement into the rich and poor halves of pots in individual species.

Fig. S4. Principal components analysis of standardized calcium and magnesium concentrations in three sample types.

Fig. S5. Relationships between Ca+Mg foraging and Ca and Mg concentrations in plant tissues.

Fig. S6. Relationships between Ca+Mg foraging (expressed as difference between NPK and Ca+Mg foraging) and Ca and Mg concentrations in biomass.

## Notes

### Competing Interest Statement

The authors have declared no competing interest.

