## supplement for "Root foraging response to gradients of calcium and magnesium, essential bivalent cations with low mobility in the soil"

**Electronic supplementary material.**

Table S1. Ordinal values (Ellenberg-type values; Chytrý et al. 2020) expressing species preference to calcium content in the soil. For these indicator values, Chytrý et al. 2020 lists two variants, one with values assigned to all taxa and the other in which values for generalist taxa are replaced by “x”."

| Name | All taxa assigned values | Generalist taxa not assigned values |
|---|---|---|
| *Nardus stricta* | 2 | 2 |
| *Anthoxanthum odoratum* agg. | 3 | 3 |
| *Agrostis capillaris* | 4 | 4 |
| *Hypericum maculatum* | 4 | 4 |
| *Luzula multiflora* | 4 | 4 |
| *Crepis mollis* | 5 | 5 |
| *Festuca rubra* | 5 | x |
| *Achillea millefolium* agg. | 6 | x |
| *Alchemilla sp.* | 6 | 6 |
| *Lathyrus pratensis* | 6 | 6 |
| *Leontodon hispidus* | 6 | 6 |
| *Plantago lanceolata* | 6 | x |
| *Ranunculus acris* | 6 | x |
| *Trifolium repens* | 6 | 6 |
| *Trisetum flavescens* | 6 | x |
| *Trifolium pratense* | 7 | x |

Tab S2. Composition of classical and modified 20% Knop solution (mg/l). Note the absence of Ca and Mg in the modified solution. The modified solution was used in the experiment.

| | classical | modified |
|---|---|---|
| $K_2SO_4$ | | 36 |
| $MgSO_4.7H_2O$ | 50 | |
| $KH_2PO_4$ | 50 | 50 |
| $NH_4NO_3$ | | 68 |
| $Ca(NO_3)_2.4H_2O$ | 200 | |
| KCl | 25 | 25 |
| $KNO_3$ | 50 | 50 |
| $FeCl_3.6H_2O$ | 5 | 5 |
| **Total N** | **30.66** | **30.73** |
| **Total S** | **6.5** | **6.93** |

Table S3. Tests of differences in Ca and Mg concentrations in experimental pots. There were four samplers arranged along a line in each tested pot, going from the dolomite-enriched patch to the opposite end of the pot. Mixed model analysis of variance with pot as a random factor, with random intercept and random effects of position and date. Tests were done with Type III sum of squares and Satterthwaite's approximation of denominator degrees of freedom. Models were fitted using lmer function from the lme4 package and tested by lmerTest package.

Calcium

| | Sum Sq | Mean Sq | NumDF | DenDF | F value | Pr(>F) | |
|---|---|---|---|---|---|---|---|
| species | 490.2 | 35.0 | 14 | 32.477 | 0.8400 | 0.6241 | |
| **position** | **19768.1** | **6589.4** | **3** | **78.251** | **158.0758** | **< 2.2e-16** | ******* |
| **date** | **3837.9** | **1919.0** | **2** | **45.126** | **46.0353** | **1.270e-11** | ******* |
| species:position | 2403.5 | 57.2 | 42 | 73.460 | 1.3728 | 0.1165 | |
| species:date | 1447.1 | 51.7 | 28 | 40.116 | 1.2398 | 0.2622 | |
| **position:date** | **3454.8** | **575.8** | **6** | **210.315** | **13.8134** | **3.202e-13** | ******* |

Magnesium

| | Sum Sq | Mean Sq | NumDF | DenDF | F value | Pr(>F) | |
|---|---|---|---|---|---|---|---|
| species | 42.19 | 3.013 | 14 | 32.597 | 1.5729 | 0.14052 | |
| **position** | **859.19** | **286.396** | **3** | **66.271** | **149.5011** | **< 2.2e-16** | ******* |
| **date** | **169.26** | **84.628** | **2** | **48.097** | **44.1766** | **1.287e-11** | ******* |
| species:position | 107.54 | 2.560 | 42 | 63.118 | 1.3365 | 0.14633 | |
| species:date | 85.95 | 3.070 | 28 | 43.379 | 1.6023 | 0.07968 | . |
| **position:date** | **61.42** | **10.236** | **6** | **212.244** | **5.3435** | **3.704e-05** | ******* |

Table S4. Average daily temperature. average temperature at 7 AM, 12 AM and 7 PM Průhonice during the experiment (March-September 2022).

| | March | April | May | June | July | August | September |
|---|---|---|---|---|---|---|---|
| Average temperature (C°) | 3.8 | 7.5 | 16.1 | 20.6 | 20.7 | 21.0 | 13.3 |
| Average temperature at 7 AM | -1.4 | 5.7 | 14.6 | 18.3 | 17.5 | 17.8 | 11.2 |
| Average temperature at 12 | 10.4 | 11.7 | 21.2 | 25.2 | 24.3 | 25.2 | 17.6 |
| Average temperature at 7 PM | 3.1 | 6.3 | 14.2 | 19.4 | 20.5 | 20.5 | 12.1 |

Table S5. Correlation coefficients of Ca and Mg content in plant biomass across individual sample types. Correlations are calculated at the level of species, i.e. n = 16.

| | Ca-Experiment aboveground | Ca-Experiment root control | Ca-Field aboveground | Mg-Experiment aboveground | Mg-Experiment root control | Mg-Field aboveground |
|---|---|---|---|---|---|---|
| Ca-Experiment aboveground | 1 | 0.568 | 0.767 | 0.514 | 0.71 | 0.671 |
| Ca-Experiment root control | 0.568 | 1 | 0.508 | 0.854 | 0.905 | 0.421 |
| Ca-Field aboveground | 0.767 | 0.508 | 1 | 0.496 | 0.645 | 0.961 |
| Mg-Experiment aboveground | 0.514 | 0.854 | 0.496 | 1 | 0.758 | 0.398 |
| Mg-Experiment root control | 0.71 | 0.905 | 0.645 | 0.758 | 1 | 0.537 |
| Mg-Field aboveground | 0.671 | 0.421 | 0.961 | 0.398 | 0.537 | 1 |

Table S6. Analysis of variance of Ca and Mg content in biomass across species and across sample types (aboveground biomass from the field, aboveground biomass from the foraging experiment, and root biomass from the poor (control) half of the pot in the foraging experiment). Response variables have been log-transformed for the analysis.

Calcium

| | Df | Sum Sq | Mean Sq | F value | Pr(>F) | |
|---|---|---|---|---|---|---|
| species | 15 | 28.1798 | 1.8787 | 51.4849 | < 2.2e-16 | *** |
| sampletype | 2 | 23.5452 | 11.7726 | 322.6306 | < 2.2e-16 | *** |
| interaction | 30 | 8.4036 | 0.2801 | 7.6768 | 7.846e-15 | *** |
| Residuals | 96 | 3.5030 | 0.0365 | | | |

Magnesium

| | Df | Sum Sq | Mean Sq | F value | Pr(>F) | |
|---|---|---|---|---|---|---|
| species | 15 | 30.363 | 2.0242 | 68.481 | < 2.2e-16 | *** |
| sampletype | 2 | 53.039 | 26.5196 | 897.190 | < 2.2e-16 | *** |
| interaction | 30 | 11.671 | 0.3890 | 13.162 | < 2.2e-16 | *** |
| Residuals | 96 | 2.838 | 0.0296 | | | |

Table S7. Tests of differences in Ca and Mg content in biomass between grasses-forbs and among sample types (same as in S5). Response variables have been log-transformed for the analysis. Mixed model analysis of variance with species as a random factor. Tests were done with Type III sum of squares and Satterthwaite's approximation of denominator degrees of freedom. Models were fitted using lmer function from the lme4 package and tested by lmerTest package.

Calcium

| | Sum Sq | Mean Sq | NumDF | DenDF | F value | Pr(>F) | |
|---|---|---|---|---|---|---|---|
| grass vs. forb | 3.1296 | 3.1296 | 1 | 13.961 | 36.7421 | 2.964e-05 | *** |
| sampletype | 16.7490 | 8.3745 | 2 | 124.012 | 98.3170 | < 2.2e-16 | *** |
| interaction | 1.3614 | 0.6807 | 2 | 124.012 | 7.9917 | 0.0005437 | *** |

Magnesium

| | Sum Sq | Mean Sq | NumDF | DenDF | F value | Pr(>F) | |
|---|---|---|---|---|---|---|---|
| grass vs. forb | 2.733 | 2.7328 | 1 | 13.906 | 24.0314 | 0.0002379 | *** |
| sampletype | 44.293 | 22.1466 | 2 | 123.956 | 194.7522 | < 2.2e-16 | *** |
| interaction | 0.418 | 0.2089 | 2 | 123.956 | 1.8369 | 0.1636233 | |

Fig. S1: Calcium and magnesium concentrations in the pots during the experiment. I, II II are sampling dates (early March, mid-April and before the harvest towards the end of May), red line average field concentrations of Ca and Mg,  A to D are positions, with A being the position within the Ca+Mg rich patch, and D being the position most distant from it (see Fig. 1 in the main text). Error bars are 95% confidence intervals. For the tests and differences among species see the Table S_FRITY.

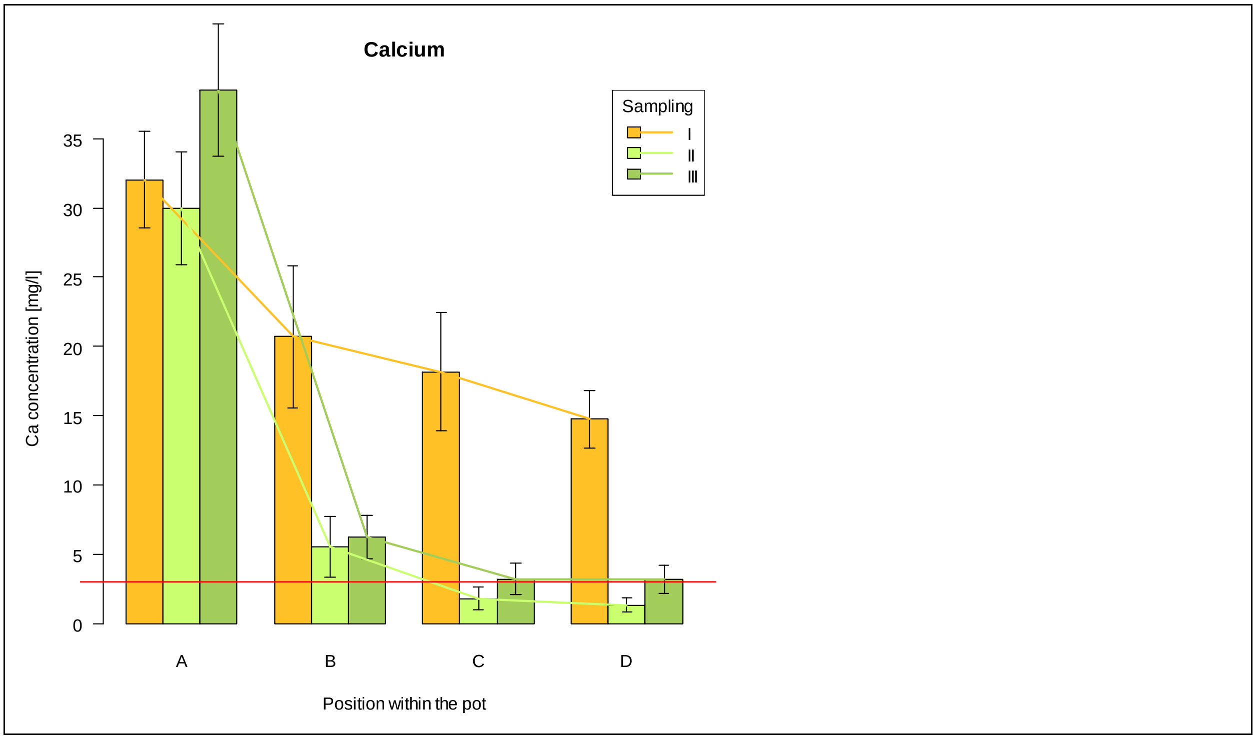

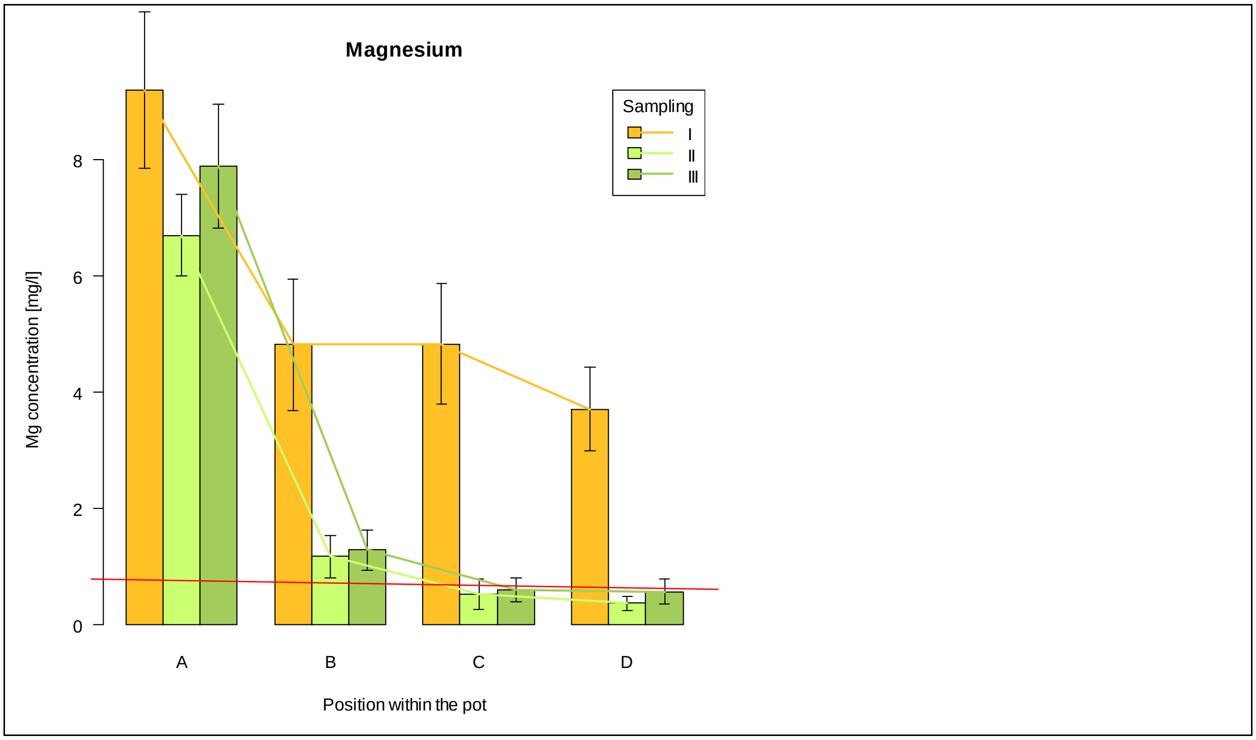
Magnesium
Sampling
I
II
III
Mg concentration [mg/l]
0
2
4
6
8
A
B
C
D
Position within the pot

Fig. S2. Experimental plants in the greenhouse at the harvest time; the patch centres are marked by coloured plastic sticks.

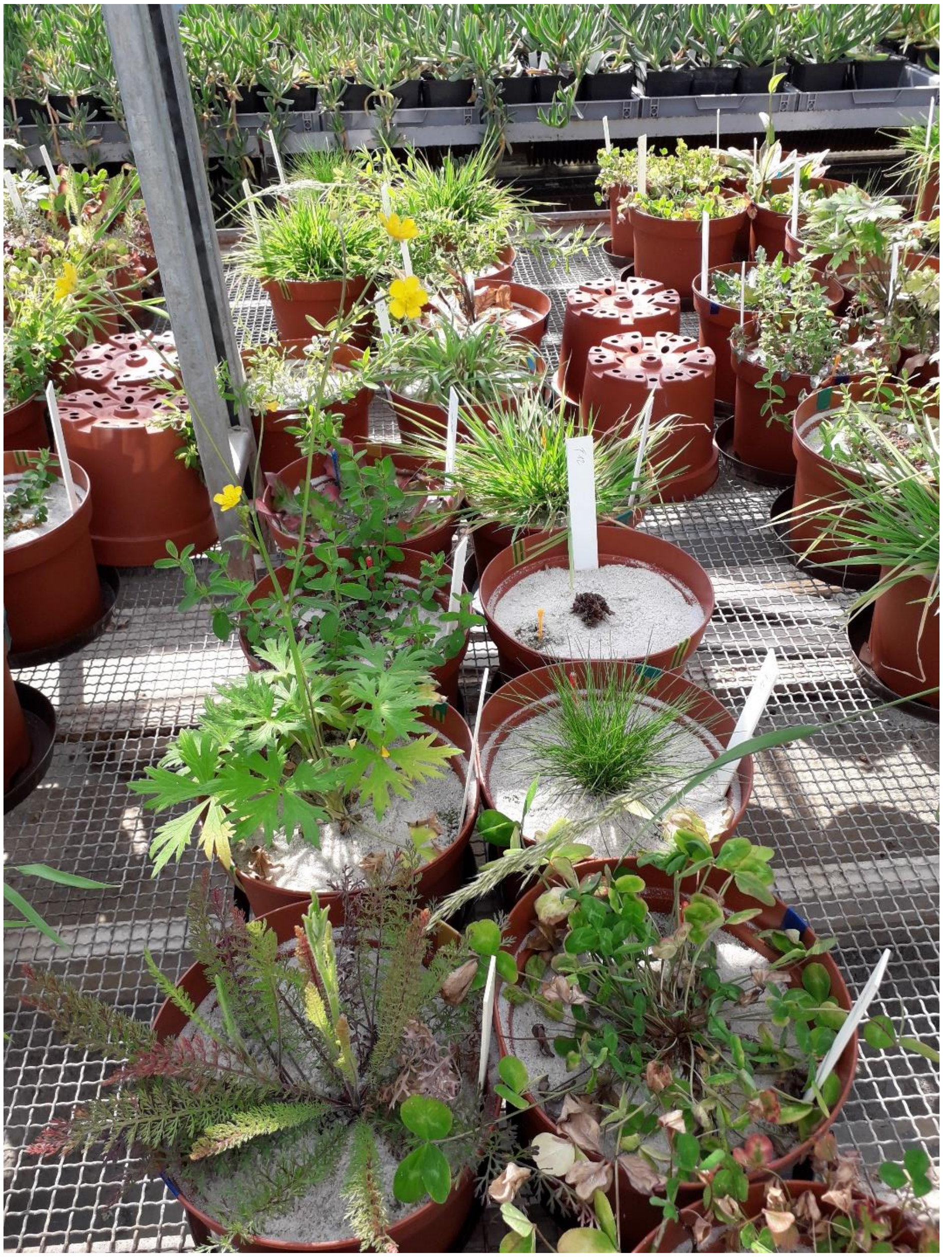

Fig. S3 Root placement into rich and poor halves in individual species. Each point indicates one pot. The lines were fitted using standard major axis regression forced through origin. The blue line is the standard major axis regression line, the dashed lines show 95% confidence intervals for the slope parameter; the black line is the y=x line (expectation under the assumption of no preferential placement). Foraging is considered significant if the slope is greater than one and the 95% confidence interval of it does not cover one.

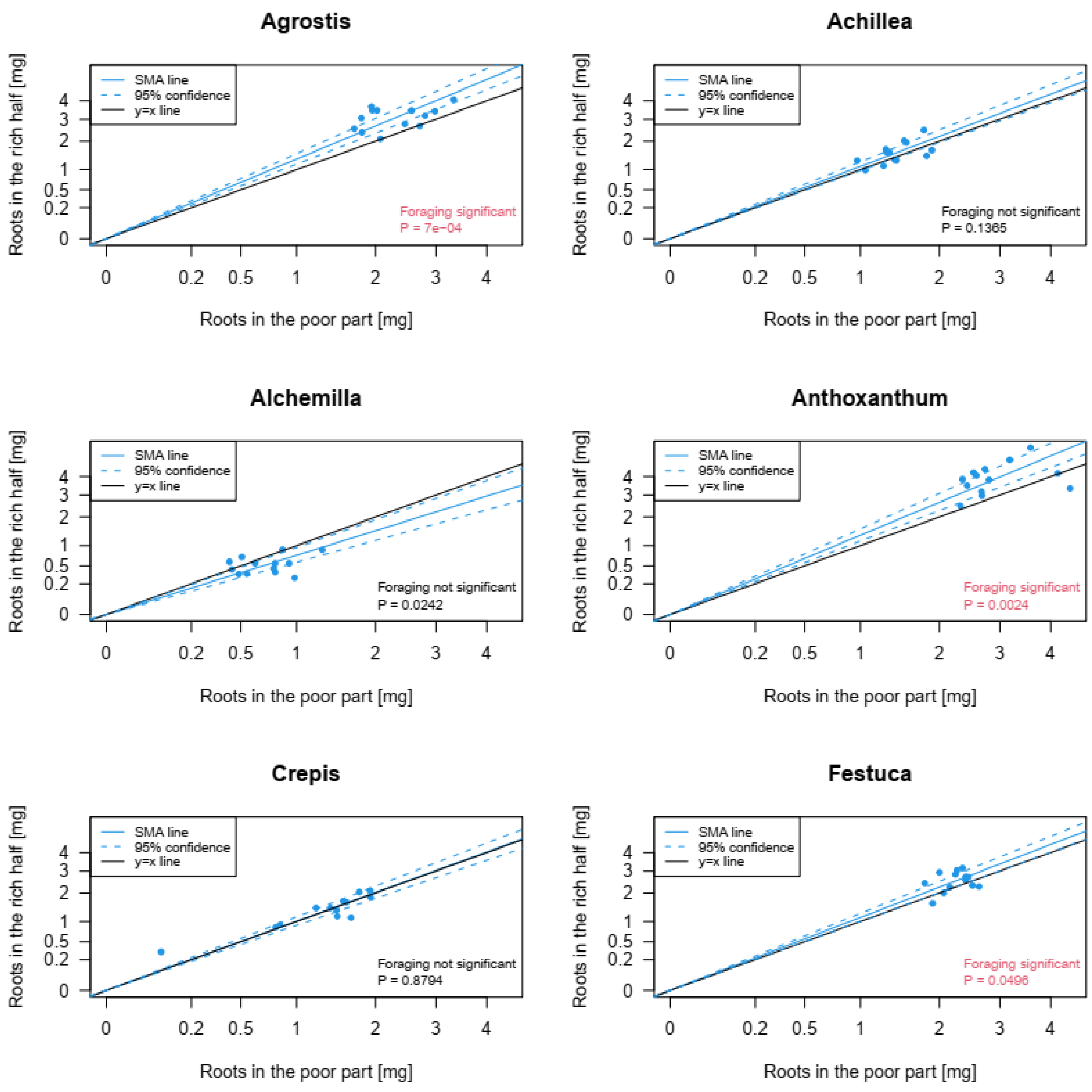

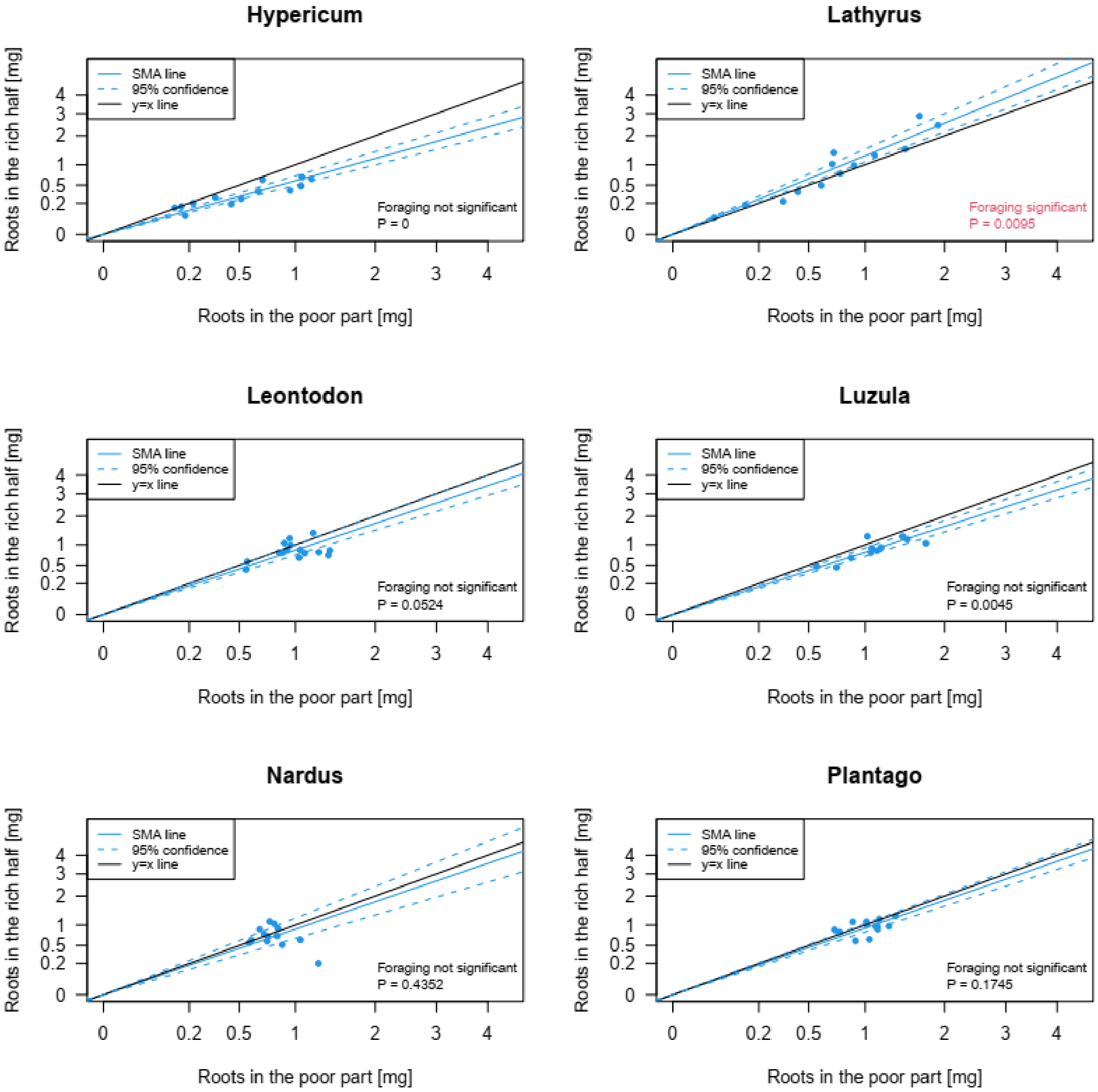
Hypericum
SMA line
95% confidence
y=x line
Foraging not significant
P = 0
Roots in the rich half [mg]
Roots in the poor part [mg]
Lathyrus
SMA line
95% confidence
y=x line
Foraging significant
P = 0.0095
Roots in the rich half [mg]
Roots in the poor part [mg]
Leontodon
SMA line
95% confidence
y=x line
Foraging not significant
P = 0.0524
Roots in the rich half [mg]
Roots in the poor part [mg]
Luzula
SMA line
95% confidence
y=x line
Foraging not significant
P = 0.0045
Roots in the rich half [mg]
Roots in the poor part [mg]
Nardus
SMA line
95% confidence
y=x line
Foraging not significant
P = 0.4352
Roots in the rich half [mg]
Roots in the poor part [mg]
Plantago
SMA line
95% confidence
y=x line
Foraging not significant
P = 0.1745
Roots in the rich half [mg]
Roots in the poor part [mg]

Fig. S4. Principal components analysis of standardized Ca and Mg concentrations in three sample types. Values have been averaged over all samples from one sample types and one species. Exp-AG – aboveground biomass from the experiment, Exp-R – root biomass from the experiment, Field-AG – aboveground biomass from the field. Numbers in parentheses are proportions of variation explained by individual axes.

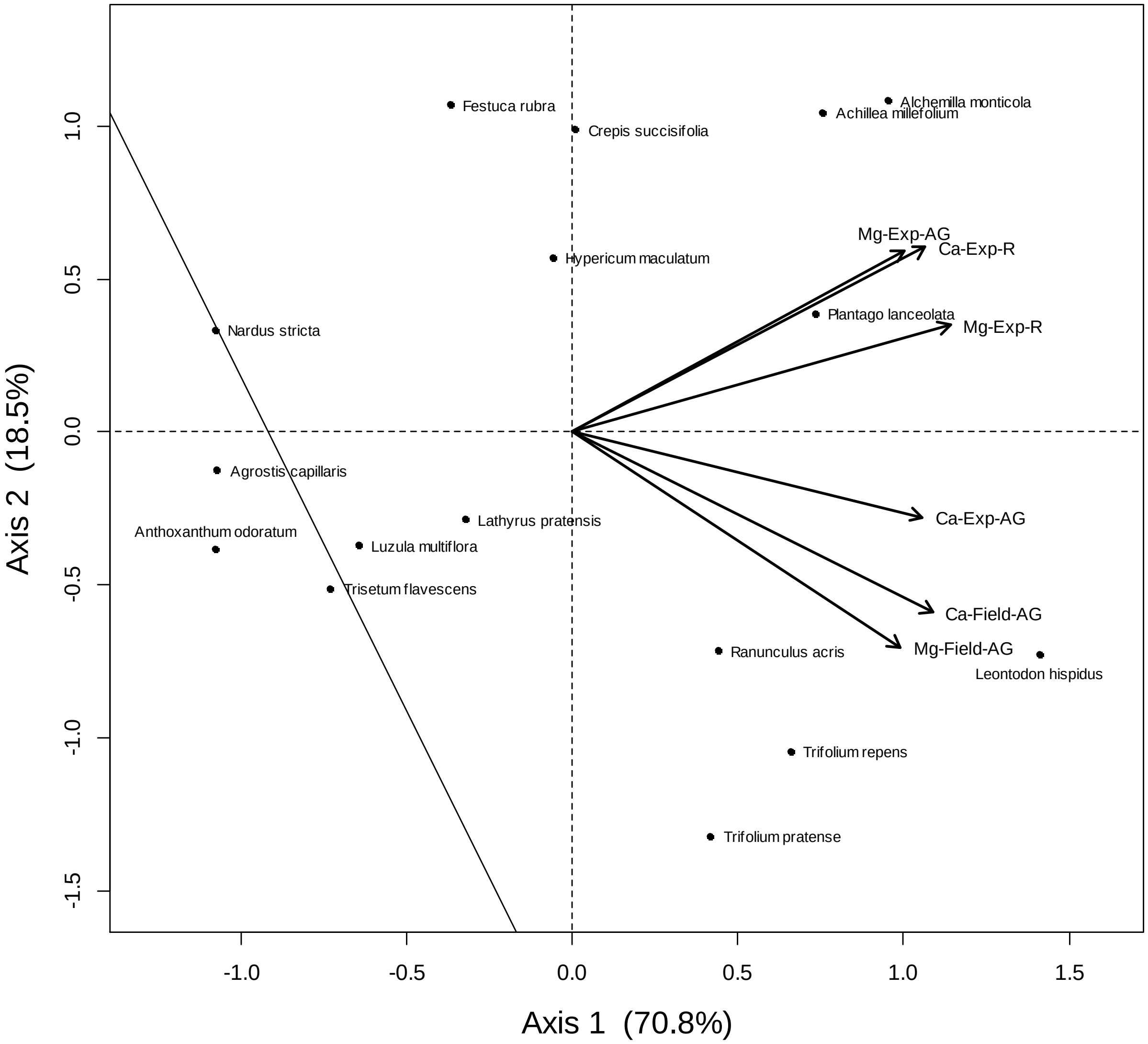

Fig. S5. Relationships between Ca+Mg foraging and Ca and Mg concentrations in biomass. Ca and Mg shoot concentration is the concentration of both elements in aboveground shoots (both in experiment and in the field), Ca concentration is concentration of calcium in all three sample types, Mg concentration is concentration of calcium in all three sample types.

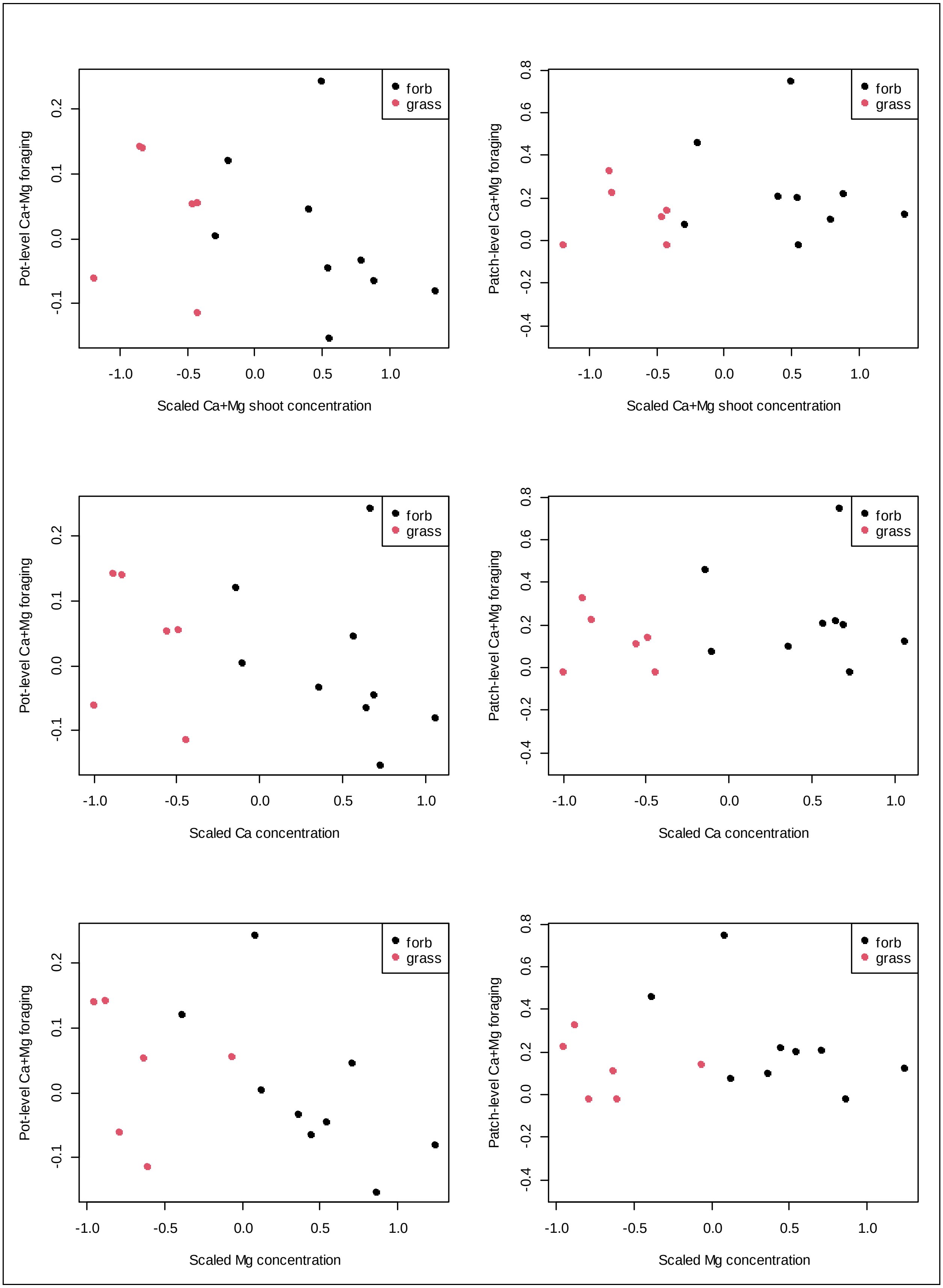

forb
grass
Pot-level Ca+Mg foraging
Patch-level Ca+Mg foraging
Scaled Ca+Mg shoot concentration
Scaled Ca concentration
Scaled Mg concentration

Fig. S6. Relationships between Ca+Mg foraging and Ca and Mg concentrations in biomass. Foraging is expressed as difference between NPK and Ca+Mg foraging; negative values thus mean that Ca+Mg foraging in the given species is weaker relative to NPK foraging. Ca+Mg shoot concentration is the concentration of both elements in aboveground shoots (both in experiment and in the field), Ca concentration is concentration of calcium in all three sample types, Mg concentration is concentration of magnesium in all three sample types. In all analyses, effect of concentration on foraging was significant at $\alpha=0.05$ (except for patch-level foraging and Ca concentration, which was only marginally significant); grass/nongrass was significant only for shoot concentrations.

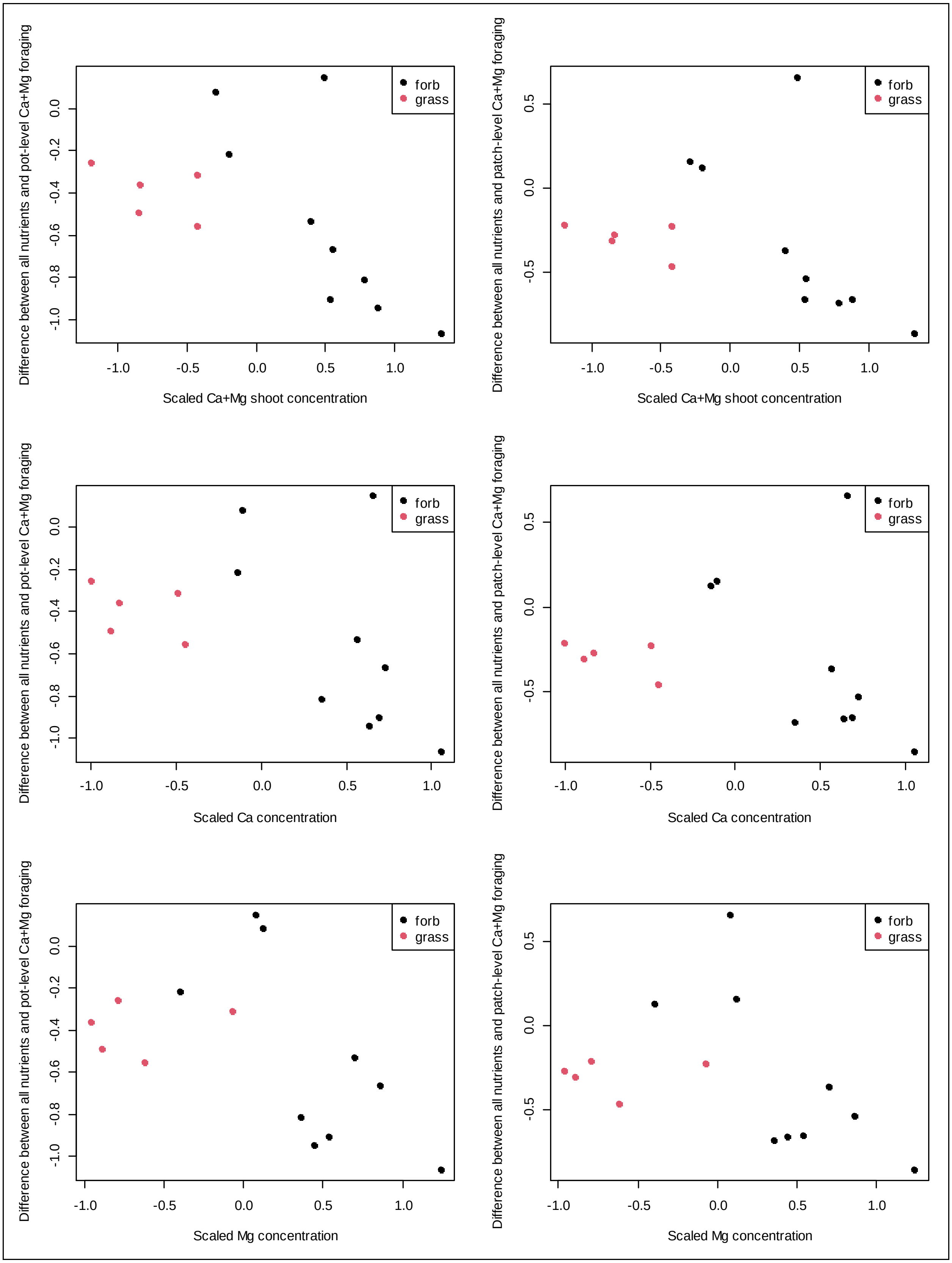
Difference between all nutrients and pot-level Ca+Mg foraging
Scaled Ca+Mg shoot concentration
forb
grass
Difference between all nutrients and patch-level Ca+Mg foraging
Scaled Ca+Mg shoot concentration
forb
grass
Difference between all nutrients and pot-level Ca+Mg foraging
Scaled Ca concentration
forb
grass
Difference between all nutrients and patch-level Ca+Mg foraging
Scaled Ca concentration
forb
grass
Difference between all nutrients and pot-level Ca+Mg foraging
Scaled Mg concentration
forb
grass
Difference between all nutrients and patch-level Ca+Mg foraging
Scaled Mg concentration
forb
grass
